# How neural circuits achieve and use stable dynamics

**DOI:** 10.1101/668152

**Authors:** Leo Kozachkov, Mikael Lundqvist, Jean-Jacques Slotine, Earl K. Miller

## Abstract

The brain consists of many interconnected networks with time-varying activity. There are multiple sources of noise and variation yet activity has to eventually converge to a stable state for its computations to make sense. We approached this from a control-theory perspective by applying contraction analysis to recurrent neural networks. This allowed us to find mechanisms for achieving stability in multiple connected networks with biologically realistic dynamics, including synaptic plasticity and time-varying inputs. These mechanisms included anti-Hebbian plasticity, synaptic sparsity and excitatory-inhibitory balance. We leveraged these findings to construct networks that could perform functionally relevant computations in the presence of noise and disturbance. Our work provides a blueprint for how to construct stable plastic and distributed networks.

## 2 Introduction

The brain is comprised of networks that are highly dynamic and noisy. Neural activity fluctuates from moment to moment and varies considerably between experimentally identical trials (Latimer et al., 2015; Lundqvist et al., 2016; 2018; Churchland et al., 2011). These fluctuations can be due to a variety of factors including variability in membrane potentials, inputs, plastic changes due to recent experience and so on. Yet, in spite of these fluctuations, networks must achieve computational stability. Despite being “knocked around” by different starting conditions and noise, networks must reach a highly consistent state for their computations to make sense.

The mechanisms that produce neural network stability have been characterized primarily in recurrent neural networks (RNNs)--a general form of brain network—in cases where the network weights are fixed and the input the network receives is constant (Fang and Kincaid 1996; Dayan and Abbot 2005). These stability conditions are bounds on the eigenvalues of the weight matrix and prevent networks from “blowing up”, that is, from running away to high levels of excitation (Fang and Kincaid 1996; Matsuoka 1992). This is an important finding but it is not the whole story. Eigenvalue analysis of the weight matrix is only guaranteed to work in RNNs receiving constant input and with fixed synaptic weights (or weights that change very slowly). Biological networks, however, have plastic synaptic weights that change rapidly under constant bombardment from environmental inputs.

Such “dynamic stability” can be studied using contraction analysis, a concept developed in control theory. Unlike a chaotic system where perturbations and distortions can be amplified over time, the population activity of a contracting network will converge towards the same trajectory, thus achieving stable dynamics (Figure 1). One way to understand contraction is to represent the state of a network at a given time as a point in the network’s ‘state-space’. A commonly used state-space in neuroscience is the space spanned by the possible firing rates of all the networks’ neurons. A particular pattern of neural firing rates corresponds to a point in this state-space. As the activity of each neuron changes, this point moves around and traces out a particular trajectory. In a contracting network, all such trajectories converge.

**Figure 1:**
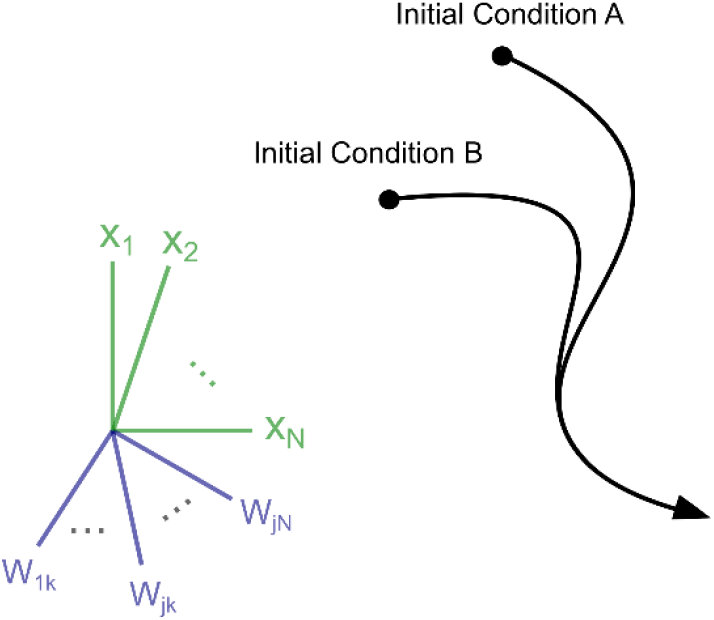
Cartoon demonstrating the contraction property. In a network with *N* neural units and *S* dynamic synaptic weights, the network activity can be described a trajectory over time in an (*N* + *S*)-dimensional space. In a contracting system all such trajectories will converge exponentially towards each other over time, regardless of initial conditions. In other words, the distance between any two trajectories shrinks to zero.

To examine how dynamic stability can be achieved with contraction under biologically realistic assumptions, we used RNNs that received time-varying inputs and had synapses that changed on biologically relevant timescales (Orhan and Ma 2019; Mongillo, Barak, and Tsodyks 2008; Lundqvist, Herman, and Lansner 2011). This revealed several classes of synaptic plasticity that naturally produced contraction, including anti-Hebbian plasticity and sparse connectivity. Further, stability is an emergent property, in the sense that two or more contracting systems can become chaotic when they interact (Ashby 2013; Lohmiller and Slotine 1998). Therefore, we also studied principles for connecting multiple networks in a way that preserved contraction as well as the functionality of each network. We then used these findings in plastic RNNs to examine how networks can perform functionally relevant computations in the presence of noise and disturbance. These computations included context-dependent sensory integration and retaining stimuli in working memory. Thus, we uncovered principles for achieving and maintaining stability in complex, modular and plastic networks.

## 3 Results

We used two main quantitative tools to characterize contraction. One is the contraction *rate*, indicating how fast trajectories reconvene following a perturbation. Another is a network’s Jacobian. The Jacobian of a dynamical system is a matrix essentially describing the local ‘traffic laws’ of nearby trajectories of the system in its state space. More formally, it is the matrix of partial derivatives describing how a change in any system variable impacts the *rate of change* of every other variable in the system. It was shown in (Lohmiller and Slotine 1998) that if the matrix measure—also known as the logarithmic norm (Söderlind 2006) – of the Jacobian is negative, then all nearby trajectories are funneled towards one another (see S.I 1.2 for technical details) which, in turn, implies that *all* trajectories are funneled towards one another.

### 3.1 Anti-Hebbian Dynamics Produce Contraction

Anti-Hebbian plasticity is the decrease of the mutual synaptic weights if the activity of two neurons are correlated. This has been observed across many brain regions and species (Hosoya, Baccus, and Meister 2005; Enikolopov, Abbott, and Sawtell 2018). It is believed to underlie important neural computations such as decorrelation of inputs (Földiák 1990). We found that anti-Hebbian plasticity produces contraction in a broad class of neural networks. Specifically, we considered neural networks of the following form:

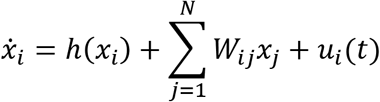

The term 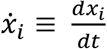 denotes the change in the activation of neuron *i* as a function of time. The term *h*(*x*_*i*_) captures the ‘self-dynamics’ of neuron *i* —the dynamics it would have in the absence of input from other neurons. The term being summed represents the weighted contribution of all the neurons in the network on the activity of neuron *i*. Finally, the term *u*_*i*_(*t*) represents external input into neuron *i*.

To ensure our results would be applicable to many different networks, we did not constrain the inputs into the RNN (except that they were not infinite), and we did not specify the particular form of *h*(*x*_*i*_) except that it be a leak term (see S.I 2.2 for what constitutes a leak term). Furthermore, we made no assumptions regarding the relative timescales of synaptic and neural activity—synaptic dynamics were treated on an equal footing as neural dynamics. In particular, let *x*_*i*_ be the activity of neuron *i*, and let *W*_*ij*_ denote the weight between neurons *i* and *j*, we considered anti-Hebbian synaptic plasticity of the following form:

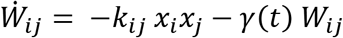

where the term *k*_*ij*_ > 0 is the anti-Hebbian plasticity learning rate for each synapse and *γ*(*t*) > 0 is a decay factor (the rate of forgetting) for each synapse. For technical reasons outlined in the supplementary, we restricted **K**, the matrix containing the *k*_*ij*_ terms, to be positive-semidefinite, symmetric, and have positive entries. A particular example of **K** satisfying these constraints is to have the learning rates of all synapses to be equal (i.e. *k*_*ij*_ = *k* > 0). Plasticity of this form produced contracting neural and synaptic dynamics, regardless of the initial values of the weights and neural activity (Figure 2 and Figure 3). In particular, we found that even if an RNN is initially not contracting, it will become contracting when subject to anti-Hebbian plasticity (Figure 3). The red trace of Figure 3.a shows that this is not simply due to the weights decaying to 0. Thus, anti-Hebbian plasticity is not only contraction preserving, it is contracting *ensuring*.

**Figure 2:**
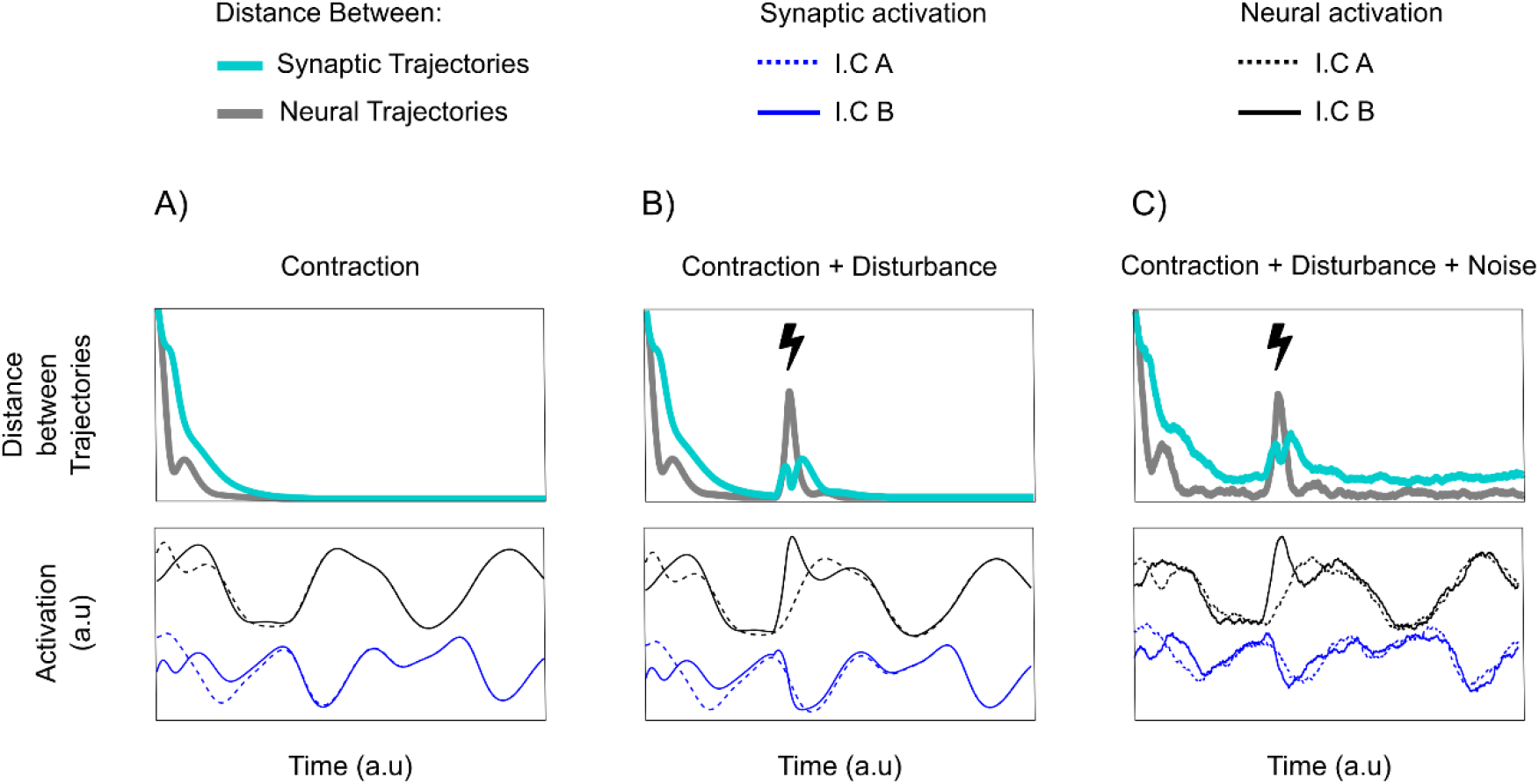
Contracting dynamics of neural and synaptic activity. Euclidean distances between synaptic and neural trajectories demonstrate exponential shrinkage over time. The top row of panels shows the distance in synaptic (teal) and activity (grey) space across simulations with distinct, randomized starting conditions. The bottom row shows the activation of a randomly selected neural unit (black) and synapse (blue) across two simulations (dotted and solid line). Panel A): Simulations of a contracting system where only starting conditions differ over simulations. B): the same as in A) but with an additional random pulse perturbation in one of the two simulations indicated by a lightning bolt symbol. C): the same as B) but with additional sustained noise, unique to each simulation.

**Figure 3:**
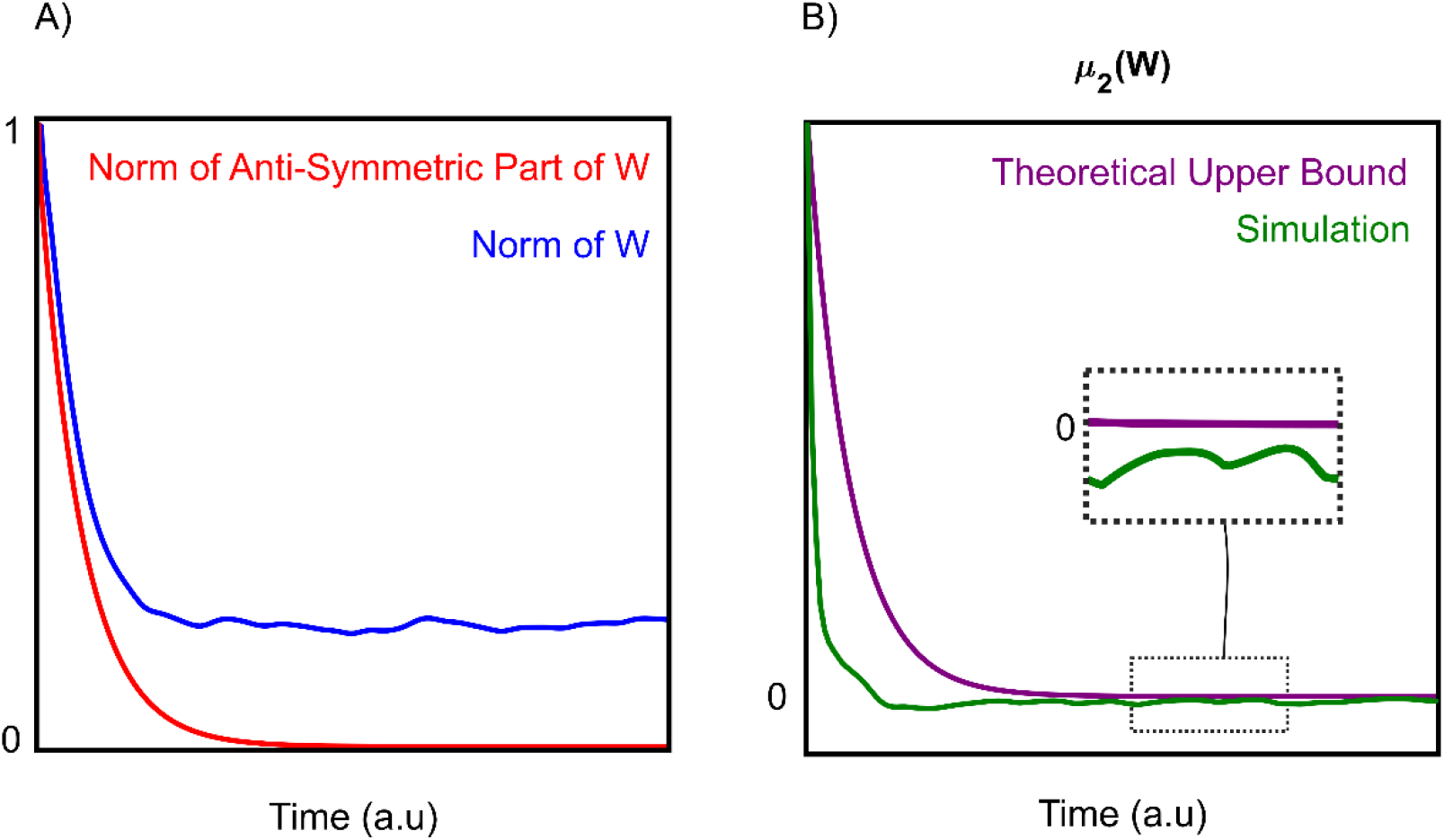
(A, red trace) The anti-Hebbian plasticity pushes the weight matrix towards symmetry. A) Plotted is a measure (the norm) of how asymmetric the weight matrix is. Red curve shows that this measure decays to zero, implying the weight matrix becomes symmetric. The blue trace shows the sum of squares of all the elements in the weight matrix. If this quantity does not decay to zero, it implies that not all the weights have decayed to zero. In (B), we plot the largest eigenvalue of the symmetric part of W (mu_2). A prerequisite for overall contraction in the network is that this quantity be less than or equal to the ‘leak-rate’ of the individual neurons. The purple line shows our theoretical upper bound for mu_2, and the green shows the actual value of mu_2 taken from a simulation. The purple decays exponentially to zero. Since the green line stays below the purple line, we can conclude that mu_2 is always less than the leak-rate of the neurons after some finite time.

To consider how anti-Hebbian plasticity works to produce contraction across a whole network, we needed to deal with the network in a holistic fashion, not by analyzing the dynamics of single neurons. To do so, we conceptualized RNNs with dynamic synapses as a single system formed by combining two subsystems—a neural subsystem and a synaptic subsystem. Contraction analysis of the overall system then boiled down to examining the interactions between these subsystems (Slotine 2003).

We found that anti-Hebbian plasticity works like an interface between these systems, producing several distinct effects that push networks toward contraction. First, it makes the synaptic weight matrix symmetric (Figure 3A, red trace). This means that the weight between neuron *i* to *j* is the same as *j* to *i*. We show this by using the fact that every matrix can be written as the sum of a purely symmetric matrix and a purely anti-symmetric matrix. An anti-symmetric matrix is one where the *ij* element is the negative of the *ji* element (*i.e. W*_*ij*_ = −*W*_*ji*_) and all the diagonal elements are zero. We then show that anti-Hebbian plasticity shrinks the anti-symmetric part of the weight matrix to zero—implying that the weight matrix becomes symmetric. Furthermore, anti-Hebbian plasticity makes the weight matrix negative semi-definite, meaning all its eigenvalues are less than or equal to zero (Figure 3A). Mathematically, we show that the symmetry of the weight matrix ‘cancels out’ off-diagonals in the Jacobian matrix (see S.I section 3) of the overall neural-synaptic system. Loosely, off-diagonal terms in the Jacobian represent potentially destabilizing cross-talk between the two subsystems. Combined with the fact that the weight matrix becomes negative semi-definite, the cancelling out of the Jacobian off-diagonals tends to funnel network dynamics towards a common path, thus producing contraction.

### 3.2 Sparse Connectivity Pushes Networks toward Contraction

Cortical synaptic connectivity is extraordinarily sparse. In the human neocortex there are about 10,000 synapses per neuron. Given that there are about 20 billion neurons in the human neocortex, this is roughly 17 orders of magnitude fewer synaptic connections than if neocortical neurons were all-to-all connected 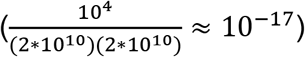. Even in local patches of cortex, such as we model here, connectivity is far from all-to-all. Our analyses revealed that sparse connectivity helps produce network contraction.

To account for the possibility that some synapses may have much slower dynamics than others, and can thus be treated as constants, we make a distinction between the total number of synapses and the total number of *dynamic* synapses. By dynamic synapse we mean a synapse whose dynamics unfold on a timescale comparable to neural dynamics. By neural dynamics we mean the change in neural activity as a function of time. A very small change in activity over a given time window would indicate a very long timescale; conversely, a very large change in activity would indicate a very short timescale. We analyzed RNNs with the structure:

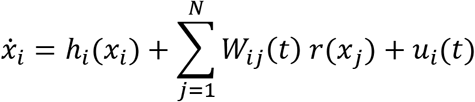

Where *h*_*i*_(*x*_*i*_) is a nonlinear leak term (see S.I section 2 for definition), and *r*(*x*_*j*_) is a nonlinear activation function. The RNNs analyzed in this section are identical to those analyzed in the previous section, with the exception of the activation term. Here we allow for a more general class of activations, whereas in the previous section we constrained *r*(*x*_*j*_) to be linear, for analytical tractability. We denote the total number of afferent synapses into neuron *i* by *p*_*i*_ and the number of afferent *dynamic* synapses by *d*_*i*_. Since the number of dynamic synapses cannot be greater than the total number of synapses, *d*_*i*_ has to be a fraction of *p*_*i*_, This means we can write it as *d*_*i*_ = *α*_*i*_*p*_*i*_, where *α*_*i*_ is a number between 0 and 1. We refer to the maximum possible absolute strength of a synapse as *W*_*max*_, the maximum possible firing rate of a neuron as *r*_*max*_ and finally the contraction rate of the *i*^*th*^ isolated neuron as *β*_*i*_. Recall from the introduction that the contraction rate measures how quickly the trajectories of a contracting system reconvene after perturbation. Under the assumption that the synapses are contracting, we show in the supplementary materials (Section 4) that if the following equation is satisfied for every neuron, then the overall network is contracting:

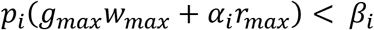

Where *g*_*max*_ is the maximum gain of any neuron in the network (see S.I section 4). Because *β*_*i*_ is a positive number, it is always possible to decrease *p*_*i*_ to the point where this equation is satisfied. Since increasing the sparsity of a network has the effect of decreasing *p*_*i*_, we may conclude that increasing the sparsity of connections pushes the system in the direction of contraction. This equation also implies that the faster the individual neurons are contracting (i.e. the larger *β* is), the denser you can connect them with other neurons while still preserving overall contraction.

### 3.3 E-I Balance Leads to Contraction in Static RNNs

Apart from making connections sparse, one way to ensure contraction is to make synaptic weights small. This can be seen for the case with static synapses by setting *α*_*i*_ = 0 in the section above. Intuitively, this is because very small weights mean that neurons cannot exert much influence on one another. If the neurons are stable before interconnection, they will remain so. Since strong synaptic weights are commonly observed in the brain, we were more interested in studying when contraction can arise irrespective of weight amplitude. Negative and positive synaptic currents are approximately balanced in biology (Mariño et al. 2005; Wehr and Zador 2003; Shu, Hasenstaub, and McCormick 2003). We reasoned that such balance might allow much larger weight amplitudes while still preserving contraction. This was indeed the case.

To show this, we studied the same RNN as in the section above, while assuming additionally that the weights are static. In particular, we show in the supplementary (section 5) that contraction can be assessed by studying the eigenvalues of the *symmetric* part of **W** (i.e. 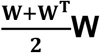). This implies the following: if excitatory to inhibitory connections are of equal amplitude (and opposite sign) as inhibitory to excitatory connections, they will not interfere with stability—regardless of amplitude (see S.I Section 5). This is because connections between inhibitory and excitatory units will be in the off-diagonal of the overall weight matrix and get cancelled out when computing the symmetric part. As an intuitive example, consider a two-neuron circuit made of one excitatory neuron and one inhibitory neuron connected recurrently (as in (Murphy and Miller 2009), Fig 1A). Assume that the overall weight matrix has the following structure:

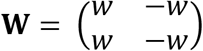

When taking that symmetric part of this matrix, the off-diagonal elements cancel out— leaving only the diagonal elements to consider. Since the eigenvalues of a diagonal matrix are simply its diagonal elements, we can conclude that if the excitatory and inhibitory subpopulations are independently contracting (*w* is less than the contraction rate of an isolated neuron), then overall contraction is guaranteed. It is straightforward to generalize this simple two-neuron example to circuits achieving E-I balance through interacting *populations* (see Supp Section 5). It is also straightforward to generalize to the case where E-I and I-E connections do not cancel out exactly neuron by neuron, but rather they cancel out in a statistical sense where the mean amplitudes are matched (Supp Section 5).

Thus far, we have described several sufficient conditions that ensure contracting dynamics in networks made of dynamic neurons and synapses. A key question is: *Can contracting dynamics be used to perform useful neural computations*? In the following sections we investigate the computational aspects of contracting networks.

### 3.4 Echo-State Networks Are Special Cases of Contracting RNNs

As can be seen in Figure 2.b, contracting systems have ‘fading memories’. This means that past events will affect the current state, but that the impact of a transient perturbation gradually decays over time. Consider the transient input in Figure 2.b (black lightning bolt) presented on only one of the two trials to the network. Because the input is only present on one trial and not the other, we call it a disturbance. Once this disturbance is presented, the distance between the trajectory corresponding to one trial and the trajectory corresponding to the other trial grows, meaning that they start to behave differently. However, after the disturbance is removed, the distance between the network’s trajectories starts shrinking back to zero again, meaning that the trajectories behave similarly.

Thus, the network does not hold onto the memory of the disturbance indefinitely—the memory fades away. A similar property has been used in Echo State Networks (ESNs) to perform useful brain-inspired computations (Jaeger 2001; Pascanu and Jaeger). These networks are an alternative to classical attractor models in which neural computations are performed by entering stable states rather than by ‘fading memories’ of external perturbations (Buonomano and Maass 2009). Because of the ‘fading memory’ property displayed by our contracting systems, we suspected that they might be related to ESNs. We investigated this next.

There are several distinctions between the networks described here and ESNs: 1) ESNs are discrete-time dynamical systems. This means that their states do not evolve continuously with time, but rather in ‘steps’. We consider continuous time networks here. While attempts have been made to find ‘Echo-State Properties’ for leaky-integrator RNNs, these have all relied on discretization of the continuous dynamics. 2) ESNs don’t have dynamic synapses and 3) The ESN ‘metric’ (which measures distances in state space) is not allowed to be time-varying. This means that the “yardstick” by which distances are measured in an ESNs state space never change, thus limiting the scope of networks classifiable as ESNs. However, by removing dynamic synapses, setting the metric we use to prove contraction equal to the identity metric, and switching to a discrete time RNN, we could derive the so-called ‘Echo state condition’ as a special case of the contracting networks considered here (see S.I section 5). It therefore follows that all the useful neural computations that have been performed by ESNs can automatically be performed by special instances of the networks considered in our work. However, by working within the framework of contraction analysis we were able to study networks both with dynamic synapses and non-stationary metrics. This allowed for greater complexity in the network dynamics while preserving the “fading memory” property. Next, we demonstrate how this additional freedom and complexity of dynamic RNNs can be applied to known problems in neuroscience.

### 3.5 Inter-areal Coupling Controls Operating Point

Neural responses to distinct stimuli or contexts should be separable from one another to enable downstream readout (Rigotti, et al., 2013). This is often determined by first averaging activity across time and trials for each experimental condition and then attempting to separate the averages linearly with hyperplanes. However, increasing evidence suggests that neural activity is highly dynamic variable from moment-to-moment and trial-to-trial (Lundqvist, et al., 2016; Wei, Inagaki, Li, Svoboda, & Druckmann, 2019; Denfield, Ecker, Shinn, Bethge, & Tolias, 2018). Therefore, it is neural *dynamics* that should be separable, not just averaged activity. The brain, after all, works in real time—not by averaging. One way to achieve context-dependent separation is by constraining the neural dynamics corresponding to a particular experimental condition to exist inside a ball of some radius around a point in state space. By moving these points—which we will call neural operating points—sufficiently far apart, one can potentially ensure that the neural dynamics do not overlap and thus ensure they are linearly separable. We therefore tested if networks consider here can guarantee linear downstream readout via contextual control of neural operating points.

There are at least two ways to control neural operating points in a contracting system: 1) By injecting tonic input; 2) By changing the network structure. Tonic input has been used in models of neural dynamics (Remington, Narain, Hosseini, & Jazayeri, 2018; Mante, Sussillo, Shenoy, & Newsome, 2013). A persistent, contextual cue (corresponding to a rule or task demand) can provide this tonic input. We observed that it shifts the neural operating point of a contracting system to a new location by shifting the “bottom” of the basin of attraction to a new location in state space (Supp Section 6). We also found that if a time-varying stimulus is then presented on top of a tonic input, the resulting neural dynamics will be contained in a sphere around the new operating point (see S.I section 6 for derivation of the radius of this sphere). This is a manifestation of the fact that a contracting system remains contracting for any (non-infinite) input.

Another way to control the neural operating point is by varying the connection strength between *coupled* contracting networks (Figure 4). We leverage the fact that it is possible for a single contracting system to connect to an arbitrary number of other contracting systems while automatically preserving contraction of the overall system (Figure 5) (Slotine 2003). Contraction is preserved but the dynamics and activity of the networks change to a degree determined by the strength of the connections between the networks. Thus, changing the changing the degree of coupling between the networks can systemically control the neural operating point of both networks (Figure 5).

**Figure 4:**
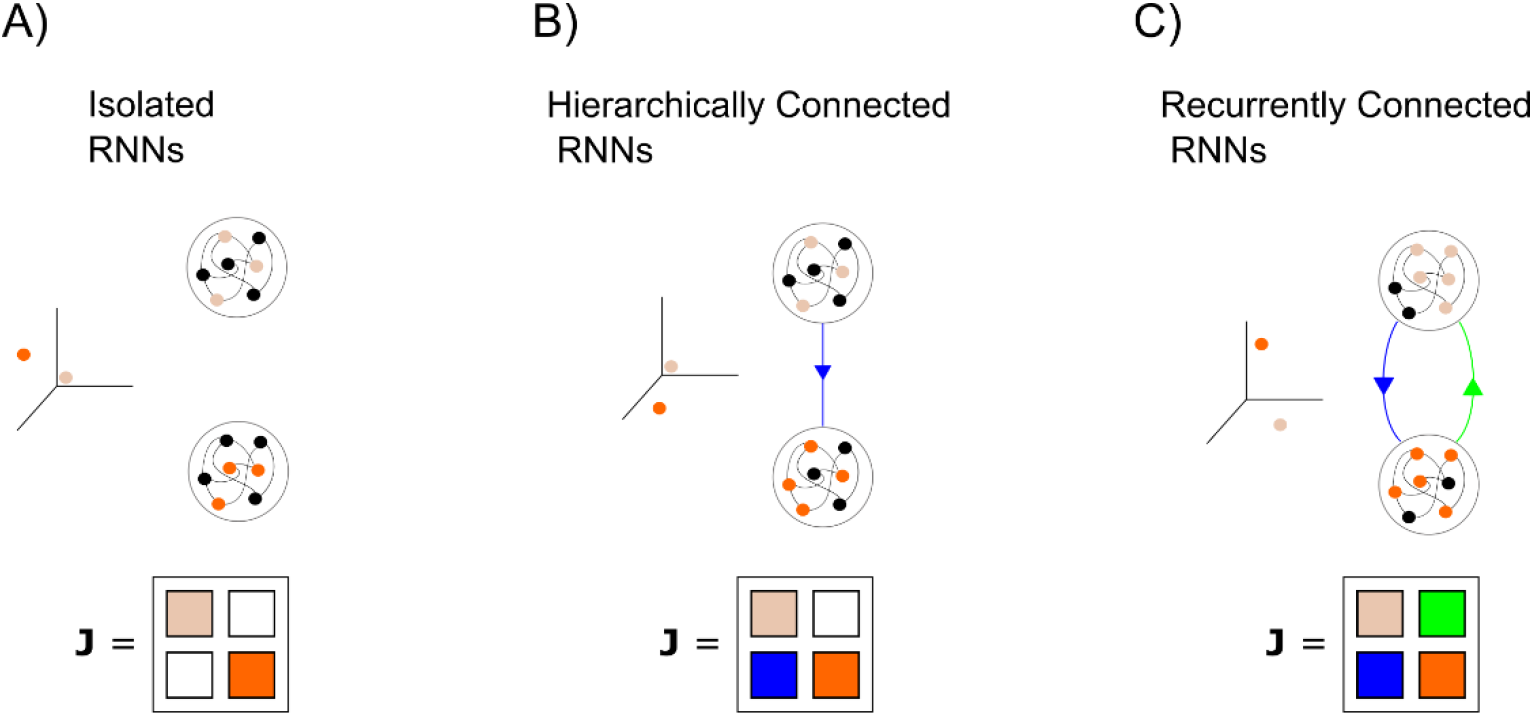
Combination properties of contracting systems. A) Two isolated, autonomous networks. The Jacobian of the overall system is block diagonal B) If one of the systems is connected to the other in a feedforward manner, the fixed point of the ‘bottom’ system will change, will the fixed point of the top system remains the same. The Jacobian of the overall system is block-triangular. C) If the systems are reciprocally connected, both systems fixed-points will change. The Jacobian is a 2 × 2 block matrix.

**Figure 5:**
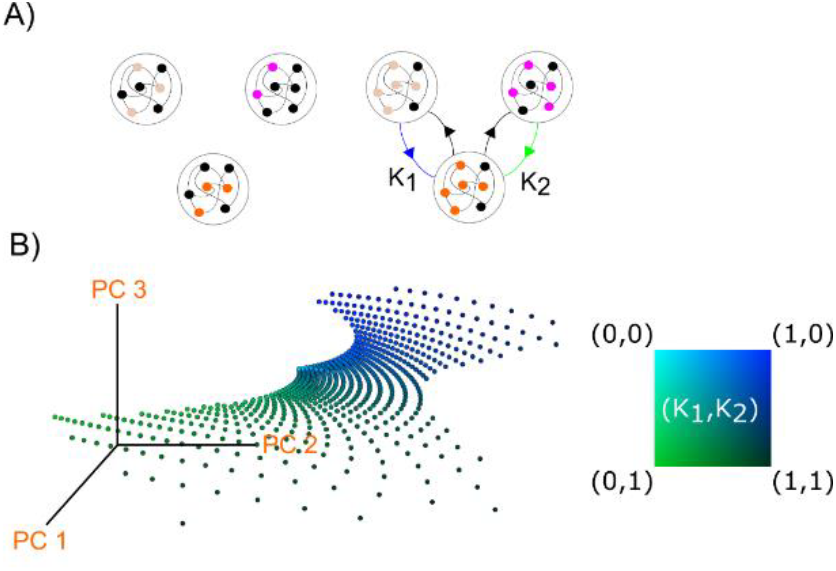
Operating point control by modulation of inter-areal connectivity. A) Left: three isolated, autonomous contracting systems. Since they are isolated, their fixed points do not depend on one another. Right: by connecting these systems, their fixed points move. B) Left: by modulating the strength of connections (k1, k2) from the two networks at the top, the fixed point of the bottom network was systematically changed. Right: the fixed points of the bottom network were plotted in space spanned by the first three principal components colored according to the value of (k1, k2).

### 3.6 Combining Contracting Networks Produces a Hierarchy of Time Constants

Elevated spiking to external stimuli is gradually prolonged as one traverses the cortical hierarchy from early sensory cortex to frontal cortex (Wasmuht, Spaak, Buschman, Miller, & Stokes, 2018; Murray J., et al., 2014). It has been suggested that shorter timescales in sensory cortex enable rapid detection of changing stimuli, while longer timescales in frontal cortex promote integration of information over time. It is not known how this hierarchical gradient is achieved. Simulations of a large-scale cortical model suggested that this is due to a gradient of increasing synaptic excitation as well as recurrent connections (Chaudhuri, Knoblauch, Gariel, Kennedy, & Wang, 2015). Here, we show instead that hierarchically combining contracting networks naturally gives rise to gradually longer time-constants of neural activity (Figure 6A). In other words, it is not strictly necessary to change the properties of the neurons to get longer time constants— it may arise from the global connectivity scheme. We therefore investigated if controlling connectivity could flexibly control the time-scale neural integration.

**Figure 6.**
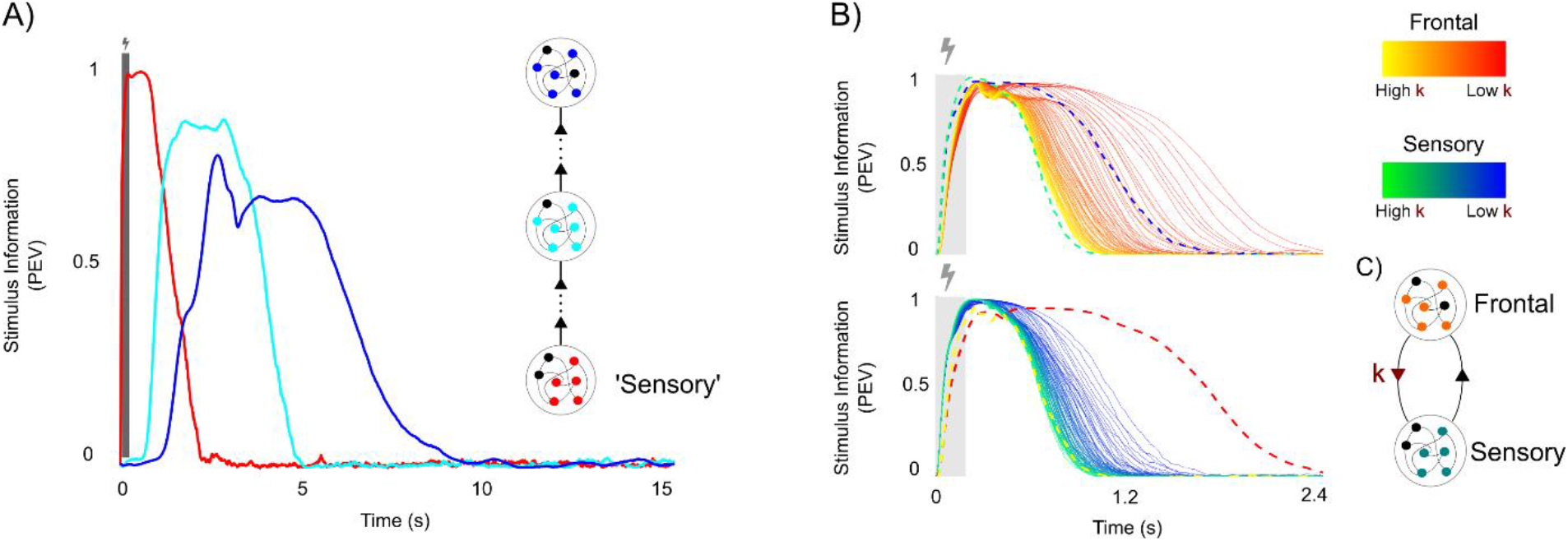
Control of time-integration by combinations of contracting systems. A) Control of integration time-constant by position in hierarchy. Hierarchical combinations of contracting systems show prolonged integration times, increasing with their position in the hierarchy. B) Modulation of top-down gain. Two networks with identical contraction rates (but different weight matrices) were reciprocally connected. The ‘sensory’ network could receive external inputs. Feedforward connections from the ‘sensory’ to the frontal network were held fixed. The top-down connections from frontal to sensory were gradually decreased in strength from k=1 towards 0, leading to a gradual increase in asymmetry in the inter-areal connectivity. For each k, external stimulus (3 different stimuli, each repeated over 100 trials) were provided to the ‘sensory’ network at t=0 (lightning bolt/grey box). Using percentage explained variance (PEV), the average time-course of stimulus information in the units’ activity was measured in both networks as a function of the asymmetry (color coded). With reduced k, the time-scale of sensory integration was prolonged, in particular in the frontal network (the dotted yellow/red lines in the sensory plot shows the two most extreme values of the above frontal plot for comparison).

First, suppose that a number of contracting subsystems are connected hierarchically. By hierarchically, we mean that while the connections *within* a subsystem can be recurrent, the connections *between* subsystems remain strictly feedforward. Our only restriction on the feedforward connectivity is that it is upper bounded in magnitude. Denote the number of subsystems as *D*. We found that the integration time of this network can scale with *m*^*D*^, where *m* > 1, which in general grows with the strength of feedforward connectivity (see S.I section 7). Thus, even with mild feedfoward connectivity strength and a few connected networks, one can get considerable increases in integration times in the higher areas. It is important to note that our results are based on upper bounds. While the integration time of this hierarchical network *can* scale exponentially with the number of subsystems, it does not have to. In practice, we did observe considerably increased information retention (almost two orders of magnitude greater than the neural time constant) in simulations as you go higher up in the hierarchy (Figure 6A), which is in agreement with experimental observations (Murray J., et al., 2014).

The cortex, of course, also has long-range feedback projections. Thus, we also explored the relation of feedback connectivity to integration times. In particular, we considered a model of interactions between sensory and frontal cortex. Both cortical areas were modelled by a contracting network, each with the same contraction rate, that we connected reciprocally (see S.I section 7). The strength of the feedback was determined by the positive parameter *k*, and gradually varied. We measured the timescales of the two networks by briefly presenting input into the sensory network and tracking how much information (Olejnik and Algina 2003) about the stimulus was retained in the network dynamics. A similar analysis as in the strictly feedforward case (above) showed that that decreasing *k* (weakening top-down feedback) leads to longer integration (Supp section 7). This was confirmed with simulations (Figure 6B). In other words, the level of time-integration was controlled by the level of top-down feedback. Consistent with the above results, the frontal network retained stimulus information for longer than the sensory cortex network despite the two networks having the same contraction rate. Both these results together show that longer integration times emerge naturally out of connecting contracting systems. Further, the time constant of the integration can be controlled by controlling feedback.

### 3.7 Stable Working Memory via Hybrid Contracting Systems

As discussed in section 3.4, contracting networks may be thought of as having a memory that fades with a characteristic time constant *λ* (a “decay constant”). There are many cases, however, where information has to be retained over gaps in time longer than *λ* (e.g., working memory tasks where memories much be held for seconds). This can be accomplished via *hybrid* contracting systems.

A hybrid dynamical system is one that is governed by the continuous evolution of variables (i.e., the type of model discussed so far) but also includes discrete transitions in synaptic weight changes (El Rifai & Slotine, 2006). These discrete transitions of synaptic weights have to be coordinated. This could be accomplished by a threshold or an “update” signal that, for example, changes synaptic weights only at given periods of time, mimicking the effect of dopamine (Lansner et al. 2013). Here, we report that the resulting hybrid contracting system can have both stable dynamics and retain memories that outlast shorter decay constants.

Consider a contracting neural network with dynamic synapses, as outlined in section 3.1. Recall that there can be separate decay constants for synapses vs neurons. Now present an input to the system. After transients, the system settles down to a new equilibrium state different from that before the input. Imagine that the weights are frozen at this new equilibrium (or the synaptic decay is much slower than the neural decay). In other words, synaptic weights are only updated when there are inputs to the network much like the stimulus-driven dopamine-mediated “print now” signal used in prior work (Lansner, Marklund, Sikström, & Nilsson, 2013). The network with frozen weights is still contracting but the equilibrium point it contracts to is different from that of the pre-stimulus network (Figure 7). In line with experimental findings (Spaak, Watanabe, Funahashi, & Stokes, 2017; Murray J., et al., 2017), the resulting activity of neurons are highly dynamic during stimulus presentation and the beginning of the delay, but gradually slows down towards a new stable equilibrium point later in the delay.

**Figure 7:**
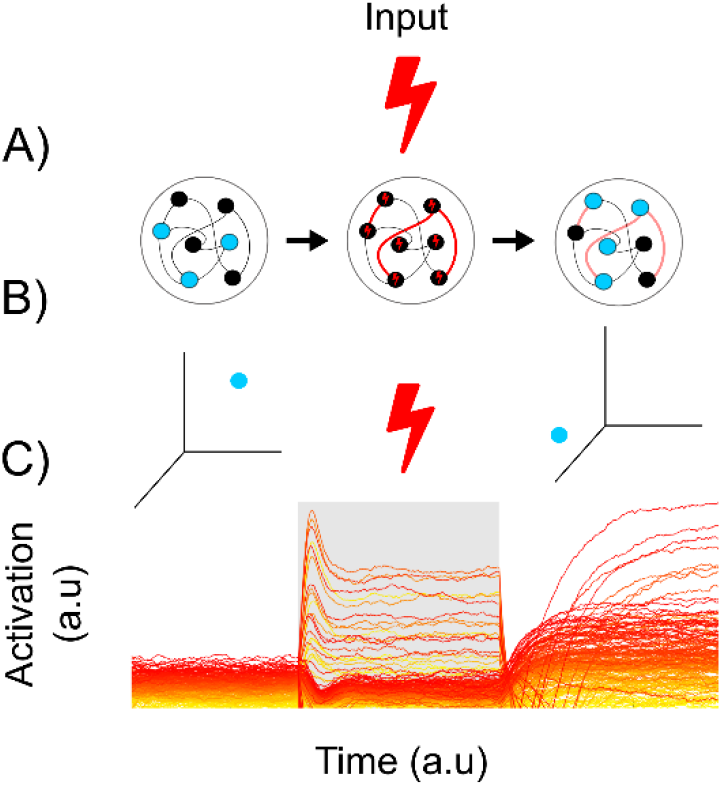
Synaptic working memory in hybrid contracting systems. The network has anti-Hebbian synaptic plasticity. (A) Left: In the absence of inputs the system has a stable fixed-point as seen in the cartoon (middle row) and neural activity (last row) sorted from most (red) to least (yellow) active unit. Middle: An input is presented to the network (grey background, lightning bolt), causing its activity to jump to a different fixed point, partly determined by the structure of the input. The synaptic weights are frozen and the input is removed. This causes the network to contract towards a new fixed-point that is informative of the now removed input.

This shows how memories in networks can outlast the neural decay constant. We show in the next section how combining this memory storage with hierarchically organized networks with increasing time constants can solve a fundamental problem of cognition: context-dependent behavior.

### 3.8 Context Dependent Behavior

Here, we construct a contracting network combining features discussed in the previous sections. We show that it can exhibit context-dependent behavior, a hallmark of cognition. Context-dependent means that behavior can change depending on the situation. We behave differently at a jazz show vs a punk show.

We combined two contracting networks: A “dynamic” network with changing synaptic weights (as discussed in section 3.7) and a “static” sensory network (Figure 8). The dynamic network was identical to the one used in Figure 7. The sensory network was set up in order to be contracting but entailed no further tweaking beyond that. We simulated the following task. At the beginning of the trial, the dynamic network was presented with one of two transient cues that instruct whether to attend to color or motion. Following a brief memory delay, the sensory network was then presented with a combined noisy color and motion stimulus and has to make a decision about the cued modality (i.e. report either the color or motion of the dots). The output of the network was a linear readout taken from the sensory network, trained to minimize the error between desired output and network output.

**Figure 8:**
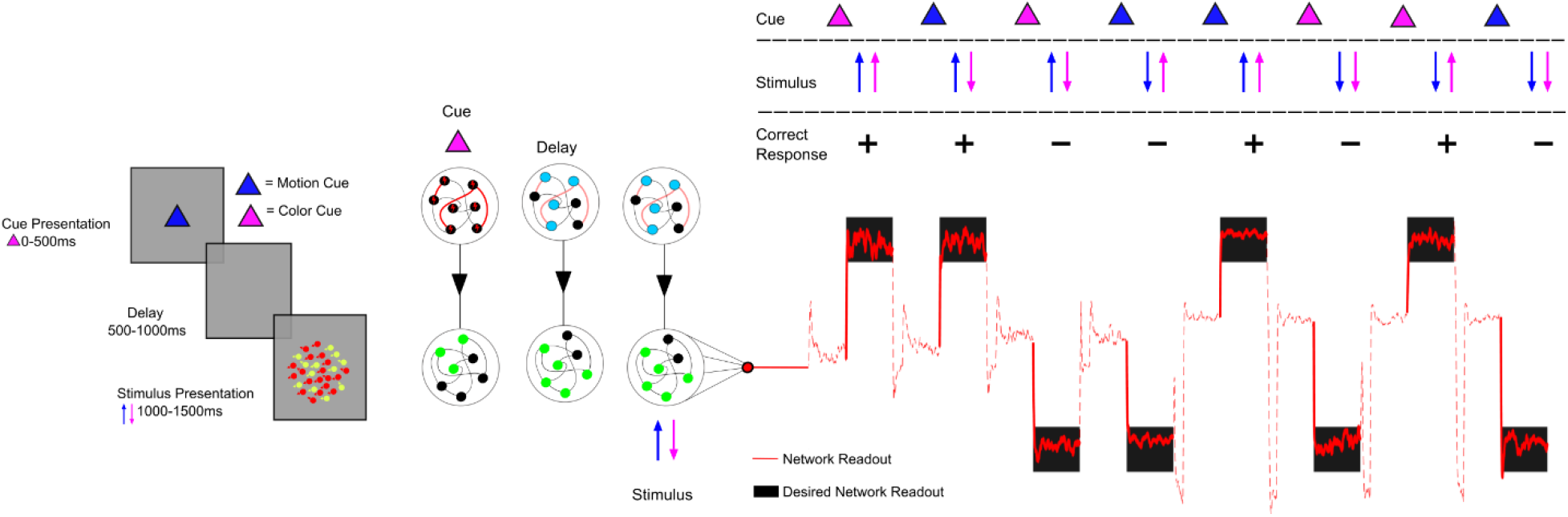
Context dependent sensory integration. A) Task design: in the task there is either a motion or color cue presented, indicating which sensory feature to pay attention to. Following a delay, sensory information is presented, and only the cued feature should dictate the response (left or right) of the network. B) Network setup: the network at the top has plastic synapses, such that it can retain the cued information, same as in Figure 6. Due to the top down connections to the sensory area the cue held in working memory provided contextual modulation. The operating point of the network thus changed with the cued context. As a result, linear read out C) could be used to make the correct response for the 8 possible trial conditions (2 cues, motion indicating left/right, color indicating left/right).

The combined networks solved the task by holding the cued modality in the network with dynamic synapses, which changed the neural operating point of the sensory network (Figure 8). This demonstrates that the properties of multiple distinct contracting systems can be combined without any fine-tuning. It also illustrates that because contracting systems have one trajectory towards which they converge, any readout (linear or nonlinear, provided that the derivative of the readout is bounded) will *also* converge (Slotine 2003). In other words, the readout is easy to read because it is linearly separable and consistent regardless of initial network conditions or noise. For this reason, we could add substantial noise to the sensory network without loss of function (Figure 8).

## 4 Discussion

We studied a fundamental question in neuroscience: how distributed neural circuits maintain stable computations in the presence of disturbance, noisy inputs and plastic change. Neurological systems have high levels of dynamical variability even between trials with identical conditions, yet produce stable behavior. We approached this problem from the perspective of dynamical systems theory, in light of the recent successes of understand neural circuits as dynamical systems (Sussillo 2014). We focused on *contracting* dynamical systems, which are yet largely unexplored in neuroscience. We did so for three reasons:

1. Contracting networks can be *input-driven*. This is important because neural circuits are typically bombarded with time-varying inputs either from the environment or from other brain areas. Previous stability analyses have focused primarily on the stability of RNNs without time-varying input. These analyses are most insightful in situations where the input into a circuit can be approximated as either absent or constant. However, naturalistic stimuli tend to be highly time-varying and complex (Steveninck et al. 1997). This allowed us to build input-driven networks that performed stable computations on time-varying inputs.
2. Contracting networks are robust to noise and disturbances. Perturbations to a contracting system are forgotten at the rate of the contraction and noise therefore does not stack up over time. Thus dynamic stability can co-exist with high trial-to-trial variability in contracting neural networks, as observed in biology.
3. Contracting networks can be combined with one another in ways that preserve contraction. This is not true of most dynamical systems which can easily ‘blow up’ when connected in feedback with one another (Ashby 2013). This combination property is important as it is increasingly clear that cognitive functions such as working memory or attention are distributed in multiple cortical and sub-cortical regions (Chatham and Badre 2015; Halassa and Kastner 2017). In particular, prefrontal cortex has been suggested as a hub that can reconfigure the cortical effective network based on task demands (Miller and Cohen 2001). Brain networks must therefore be able to effectively reconfigure themselves on a fast time-scale without loss of stability. We show how to achieve this automatically with contracting networks. Most attempts in modelling cognition, for instance working memory, tend to utilize single and often autonomous networks. Contracting networks display a combination of input-driven and autonomous dynamics, and thus have key features necessary for combining modules into flexible and distributed networks.

To understand what mechanisms lead to contraction in neural circuits, we applied contraction analysis to RNNs. For RNNs with static weights, we found that the well-known Echo State Networks are a special case of a contracting network. Since realistic synapses are complex dynamical systems in their own right, we went one step further and asked when neural circuits with dynamic synapses would be contracting. We found that anti-Hebbian plasticity and synaptic sparsity both lead to contraction in a broad class of RNNs. Anti-Hebbian plasticity exists across many brain areas and species, such as salamander and rabbit retina (Hosoya, Baccus, and Meister 2005), rat hippocampus (Lisman 1989; Kullmann and Lamsa 2007), electric fish electrosensory lobe (Enikolopov, Abbott, and Sawtell 2018) and mouse prefrontal cortex (Ruan, Saur, and Yao 2014). These dynamics can give rise to sparse neural codes which decrease correlations between neural activity and increase overall stimulus representation in the network (Földiák 1990). Because of this on-line decorrelation property, anti-Hebbian plasticity has also been implicated in predictive coding (Hosoya, Baccus, and Meister 2005; Enikolopov, Abbott, and Sawtell 2018).

For synaptic plasticity that is not necessarily anti-Hebbian, we showed (in section 3.2) that in general, synaptic sparsity pushes RNNs towards being contracting. This aligns well with the experimental observation that synaptic connectivity is typically extremely sparse in the brain. Our results suggest that sparsity may be one factor pushing the brain towards contractive behavior. It is therefore interesting that synapses are regulated by homeostatic processes where synapses neighboring an upregulated synapse are immediately downregulated (El-Boustani et al. 2018). On the same note, we also observed that balancing the connections between excitatory and inhibitory populations leads to contraction. Balance between excitatory and inhibitory inputs are often observed in biology (Mariño et al. 2005; Wehr and Zador 2003; Shu, Hasenstaub, and McCormick 2003), and could thus serve contractive stability purposes. Related computational work on spiking networks has suggested that balanced synaptic currents leads to fast response properties, efficient coding, increased robustness of function and can support complex dynamics related to movements (Denève and Machens 2016; Hennequin, Vogels, and Gerstner 2014; Lundqvist, Compte, and Lansner 2010; Brunel 2000).

We used the anti-Hebbian plasticity to build a working memory network where inputs were retained at a time-scale much longer than the contraction rate. The outcome of the plastic changes induced by a stimulus were frozen into the network and forced the network to converge towards a new trajectory unique to that input. As a result, activity was highly dynamic during input but stabilized exponentially and reached a stable plateau a few hundred millisecond later. Similar dynamics have been observed in spiking activity of recorded populations during working memory tasks in non-human primates. In addition, individual units displayed rich dynamics with time-varying selectively, as also observed experimentally (Barak, Tsodyks, and Romo 2010; Warden and Miller 2010). Earlier computational studies have also suggested a role for synaptic plasticity in working memory (Sandberg, Tegnér, and Lansner 2003; Mongillo, Barak, and Tsodyks 2008; Lundqvist, Herman, and Lansner 2011, 2012; Fiebig and Lansner 2017), but not within the framework of dynamic stability.

The combination properties of these systems allowed us to combine the functionalities of local neural circuits in simple ways to solve various simulated cognitive tasks with essentially no fine-tuning. In particular, we combined all the above properties to construct a modular network that solved a context-dependent sensory integration task. The network was noise tolerant and required no tuning, illustrating the ease with which one can build up complex functionalities from simpler ones using contracting networks.

Further, we defined the neural operating point of a contracting RNN as the point around which all its trajectories are bounded. We found that by modulating the strength of connection between combined contracting systems or by the injection of tonic input into a contracting network one could shift this operating point. This enables separation of neural trajectories. Linear separation has been discussed as an important feature of higher cognition (Rigotti et al. 2013). There is recent experimental evidence suggesting that weight matrix modulation and tonic input modulation indeed exists and may be thalamic in origin (Rikhye, Gilra, & Halassa, 2018).

We found that combining identical contracting RNNs hierarchically automatically produced a gradient of time-constants. Such gradient has been observed in cortex (Murray et al. 2014). Current models account for this phenomenon through a cortical gradient in synaptic time-constants, in other words, by imposing the gradient. We found that increasing time constants automatically occurs when connecting contracting networks into a hierarchy. This makes it broadly applicable and flexible with respect to biological detail. Furthermore, our analysis revealed that the timescales of neural computation to be regulated in a robust and stable way simply by changing the amount of inter-area top-down feedback. This opens the possibility that the integration to be controlled by cognitive processes such as attention.

Experimental neuroscience is moving in the direction of studying many interacting neural circuits simultaneously. We therefore anticipate that the presented work can provide a useful foundation for how cognition in noisy and distributed computational networks can be understood.

## Acknowledgments

We thank Pawel Herman for comments on an earlier version of this manuscript. We thank Michael Happ and all members of the Miller Lab for helpful discussions and suggestions. We thank Charles Shvartsman for code used in section 3. This work was supported by NIMH R37MH087027, ONR MURI N00014-16-1-2832, NSF 1809314, and The MIT Picower Institute Innovation Fund.

