## Supplemental Materials for "How neural circuits achieve and use stable dynamics"

**6 Setting Neural Operating Points** **29**

**7 Metric-Induced Overshoots, Longer Information Retention, and a Hierarchy of Time Constants** **30**

### 1 Preliminaries

#### 1.1 Mathematical Preliminaries

##### 1.1.1 Matrix Notions

We will denote vectors and matrices with bold font. For example: $\mathbf{x} \in \mathbb{R}^N$ is an $N$-dimensional column vector and $\mathbf{A} \in \mathbb{R}^{N \times M}$ is an $N \times M$ matrix. We use the notation $\mathbf{P} \succ 0$ to denote a positive definite matrix, which is a square matrix such that, for all $\mathbf{x} \in \mathbb{R}^N \neq \mathbf{0}$, the quadratic form $\mathbf{x}^T\mathbf{P}\mathbf{x}$ is strictly positive:

$$\mathbf{P} \succ 0 \equiv \mathbf{x}^T\mathbf{P}\mathbf{x} > 0 \tag{1.1.1}$$

When the quadratic form is non-negative (i.e $\mathbf{x}^T\mathbf{P}\mathbf{x} \geq 0$) then we say that that $\mathbf{P}$ is positive semi-definite and write $\mathbf{P} \succcurlyeq 0$. Negative definiteness and semi-definiteness are defined the same way, with the sign of the above inequalities reversed.

Every real, square matrix $\mathbf{Q}$ can be decomposed into a symmetric part and a skew symmetric part. We denote the symmetric part of $\mathbf{Q}$ as $\mathbf{Q}_s$ and define it as follows

$$\mathbf{Q}_s \equiv \frac{\mathbf{Q} + \mathbf{Q}^T}{2} \tag{1.1.2}$$

and the antisymmetric (skewsymmeric) part as

$$\mathbf{Q}_a \equiv \frac{\mathbf{Q} - \mathbf{Q}^T}{2} \tag{1.1.3}$$

Note that $\mathbf{Q} = \mathbf{Q}_s + \mathbf{Q}_a$. Also note that without ambiguity, when we say that a non-symmetric $\mathbf{Q}$ is positive definite, we mean that $\mathbf{Q}_s \succ 0$. This is because:

$$\mathbf{x}^T\mathbf{P}\mathbf{x} = \mathbf{x}^T\mathbf{P}_s\mathbf{x}$$

Substituting this equality back into (1.1.1), we see that a matrix is positive definite if and only if its symmetric part is positive definite. For more information on positive definite matrices, see section 7.1 of [3] or section 3.5.1 of [5], for example.

##### 1.1.2 Matrix Measures

We will also need the fact that a given vector norm $|\cdot|$ will induce a matrix norm $||\cdot||$

$$||\mathbf{Q}|| \equiv \sup_{|x|=1} |\mathbf{Qx}|$$

as well as a matrix measure $\mu(\cdot)$

$$\mu(\mathbf{Q}) \equiv \lim_{h\to 0} \frac{||\mathbf{I} - h\mathbf{Q}|| - 1}{h}$$

where $\mathbf{I}$ denotes the identity matrix. Different norms induce different matrix measures, but all matrix measures share certain common properties. In particular, for any matrix measure $\mu$ we have subadditivity:

$$\mu(\mathbf{X} + \mathbf{Y}) \leq \mu(\mathbf{X}) + \mu(\mathbf{Y}) \tag{1.1.4}$$

positive homogeneity:

$$\forall c \geq 0, \quad \mu(c\mathbf{X}) = c\mu(\mathbf{X}) \tag{1.1.5}$$

as well as

$$\mu(\mathbf{I}) = 1 \text{ and } \mu(-\mathbf{I}) = -1 \tag{1.1.6}$$

For more information about matrix measures the reader is referred to section 2.2.2 of [7] or section 2 of [1].In this paper we will rely primarily on the matrix measures induced by the infinity-norm $|\cdot|_\infty$ and the 2-norm $|\cdot|_2$. These norms induce the matrix measures

$$\mu_\infty(\mathbf{Q}) = \sup_i (Q_{ii} + \sum_{j\neq i} |Q_{ij}|) \tag{1.1.7}$$

$$\mu_2(\mathbf{Q}) = \lambda_{\max}(\mathbf{Q}_s) \tag{1.1.8}$$

Finally, consider the (potentially time-varying), non-singular matrix $\mathbf{\Theta}$ and the associated weighted 2-norm and weighted infinity-norm : $|\mathbf{x}|_{\Theta,2} = |\mathbf{\Theta x}|_2$. The matrix measures induced by these norms are:

$$\mu_{2,\Theta}(\mathbf{Q}) = \mu_2(\dot{\mathbf{\Theta}}\mathbf{\Theta}^{-1} + \mathbf{\Theta Q \Theta}^{-1}) \tag{1.1.9}$$

$$\mu_{\infty,\Theta}(\mathbf{Q}) = \mu_\infty(\dot{\mathbf{\Theta}}\mathbf{\Theta}^{-1} + \mathbf{\Theta Q \Theta}^{-1}) \tag{1.1.10}$$

##### 1.1.3 Matrix Vecorization

Finally, consider an arbitrary $N \times N$ matrix $\mathbf{A}$. The vectorization of $\mathbf{A}$, which we denote by $\mathbf{v}_A$, is the $N^2 \times 1$ column vector obtained by stacking the columns of $\mathbf{A}$ on top of one another, starting from the leftmost column and ending at the rightmost column. Consider the example of:

$$\mathbf{A} = \begin{bmatrix} a & b \\ c & d \end{bmatrix}$$

then

$$\mathbf{v}_A = \begin{bmatrix} a \\ c \\ b \\ d \end{bmatrix}$$

We also define the $N^2 \times N^2$ matrix $\mathbf{D}_A$ as the diagonal matrix with the elements of $\mathbf{A}$ along its diagonal, starting with the leftmost column of $\mathbf{A}$ and ending with the rightmost column. For the $\mathbf{A}$ matrix defined above, we have:

$$\mathbf{D}_A = \begin{bmatrix} a & 0 & 0 & 0 \\ 0 & c & 0 & 0 \\ 0 & 0 & b & 0 \\ 0 & 0 & 0 & d \end{bmatrix}$$

These two operators have a special relationship to the Hadamard product (element-wise matrix multiplication). Consider a matrix $\mathbf{Z}$ defined as:

$$\mathbf{Z} = \mathbf{X} \odot \mathbf{Y}$$

where $\mathbf{X}$ and $\mathbf{Y}$ are of the same size. Then we have the following relationship:

$$\mathbf{v}_Z = \mathbf{D}_X \mathbf{v}_Y \qquad (1.1.11)$$

which can be verified through direct inspection.

#### 1.2 Basic Contraction Definition

In this section we define what a contracting system is, and state several properties of these systems that will be useful for the remainder of the supplementary. The reader is referred to [4] and [6] for more details. Consider a deterministic system of the form

$$\dot{\mathbf{x}} = \mathbf{f}(\mathbf{x}, t) \qquad (1.2.1)$$

where $\mathbf{f}$ is an $N{\times}1$ nonlinear vector function and $\mathbf{x}$ is the $N{\times}1$ state vector. We assume that all quantities are real and smooth, by which we mean that any required derivative or partial derivatives exist and are continuous. Denote the $N \times N$ time-varying Jacobian of this system as $\mathbf{J}(\mathbf{x}, t) = \mathbf{J} = \frac{\partial \mathbf{f}}{\partial \mathbf{x}}$.

If there exists a matrix measure $\mu$ such that

$$\forall \mathbf{x}, t \quad \mu(\mathbf{J}) \leq -\lambda \ \text{ with } \ \lambda > 0 \qquad (1.2.2)$$

then all trajectories of (1.2.1) converge exponentially towards one another with rate $\lambda$ and the system is said to be *contracting*. In particular, let $R(t) =$

$|\mathbf{x}(t) - \mathbf{x}'(t)|$ denote the distance between any two trajectories of system (1.2.1) and assume the system is contracting. Then there exists a constant $k \geq R(0)$ such that

$$0 \leq R(t) \leq ke^{-\lambda t}$$

To avoid ambiguity in language, we now define some terms associated with the measures $\mu_1$,$\mu_2$ and $\mu_\infty$ . As in [4], consider a metric $\mathbf{M} = \mathbf{\Theta}^T\mathbf{\Theta} \succ 0$ and the following associated quantity, called the *generalized Jacobian*:

$$\mathbf{F} = \dot{\mathbf{\Theta}}\mathbf{\Theta}^{-1} + \mathbf{\Theta}\mathbf{J}\mathbf{\Theta}^{-1} \tag{1.2.3}$$

If the matrix measures induced by the 1,2, or infinity norm are negative definite when applied to $\mathbf{F}$, i.e.

$$\mu_i(\mathbf{F}) < -\lambda \;\text{ with }\; i = 1, 2, \text{ or } \infty \tag{1.2.4}$$

then we say that the system is contracting in metric $\mathbf{\Theta}$ with rate $\lambda$. Calling $\mathbf{\Theta}$ a metric is a slight abuse of language, because the real metric is $\mathbf{M} = \mathbf{\Theta}^T\mathbf{\Theta}$, but since we will always use the symbol $\mathbf{\Theta}$ in the generalized Jacobian, this won't cause any confusion.

#### 1.3 Robustness of Contracting Systems to Disturbance

Consider the system defined by (1.2.1), contracting in metric $\mathbf{\Theta}$ with rate $\lambda$. Now consider this same system, perturbed with an additive disturbance:

$$\dot{\mathbf{x}}_p = \mathbf{f}(\mathbf{x}_p, t) + \mathbf{d}(\mathbf{x}_p, t)$$

Then the distance between any trajectory of the disturbed system and the undisturbed system, denoted $R(t) = |\mathbf{x}(t) - \mathbf{x}_p(t)|$, obeys:

$$R(t) \leq \chi R(0)e^{-\lambda t} + \frac{|d|\chi}{\lambda} \tag{1.3.1}$$

where $\chi$ is an upper bound on the condition number of $\mathbf{\Theta}$ and $|d|$ is an upper bound on the norm of the disturbance.

### 2 Contracting Dynamics of A Leaky Neuron

In this section we apply contraction analysis to the dynamics of a single, leaky, input-driven neuron. The section is organized into three parts: neuron with no self-synapse, neuron with a static self-synapse, and neuron with a dynamic self-synapse. Analyzing the case of one neuron will turn out to be helpful when thinking about the case of many neurons.

#### 2.1 A Motivating Example

Consider the activation of a pure integrator neuron that is receiving constant input, $I$:

$$\dot{x} = I \tag{2.1.1}$$

This differential equation can be solved analytically, and has the solution:

$$x(t) = It + x(0) \tag{2.1.2}$$

Since the solution does not forget x(0), its initial condition, this system is not contracting. However, if we were to add a 'leak' term, for example $-x$ to (2.1.1), we would have

$$\dot{x} = -x + I \tag{2.1.3}$$

which yields the solution

$$x(t) = I + \chi e^{-t} \tag{2.1.4}$$

with $\chi = x(0) - I$. Since this system forgets $x(0)$ exponentially quickly, we may conclude that it is contracting (of course we also could have computed the

Jacobian, which in the case of (2.1.3) is $-1$). In the case of nonlinear dynamics and time-varying input, we will not be able to solve for the dynamics of $x$ explicitly to determine contracting properties–instead, we will have to analyze the Jacobian.

#### 2.2 Leaky Neuron with No Autapse

Let $x$ denote the activation of a single, isolated, leaky-integrator neuron. Assume that this neuron has no self-connection (i.e no autapse) and obeys the equation:

$$\tau \dot{x} = -\alpha x + I(t) \tag{2.2.1}$$

where $\tau > 0$ is the time-constant of integration, $\alpha > 0$ is the time-constant of the 'leak', and $I(t)$ is an external input into the neuron. If $\alpha = 0$ this would be a pure integrator neuron (see section 2.1).Dividing through by $\tau$, we may rewrite (2.2.1) as

$$\dot{x} = -\frac{\alpha}{\tau}x + \frac{1}{\tau}I(t)$$

the Jacobian of this scalar system is:

$$J = \frac{\partial \dot{x}}{\partial x} = -\frac{\alpha}{\tau} \tag{2.2.2}$$

since $\alpha$ and $\tau$ are positive, $\frac{\alpha}{\tau}$ is also positive, which means that $-\frac{\alpha}{\tau}$ is negative. Since the Jacobian is negative, we can conclude that the isolated leaky neuron is contracting for all inputs. In the paper we will consider several classes of leaky neurons, including neurons that obey the equation

$$\tau \dot{x} = h(x) + I(t) \tag{2.2.3}$$

where $h(x)$ is a nonlinear function that satisfies

$$\exists\, \beta > 0, \quad \forall\, x, \quad \forall\, t \geq 0, \quad \frac{\partial h}{\partial x} \leq -\beta < 0 \tag{2.2.4}$$

By the same arguments as above, leaky neurons of this form are also contracting in isolation. Of course, the standard leak term of $-x$ is a special case of $h(x)$. For $h(x) = -x$ we find that $\beta = 1$.

#### 2.3 Leaky Neuron with a Static Autapse

What happens if the neuron defined by (2.2.3) *does* have an autapse? In other words, assume the neural dynamics now follow

$$\tau \dot{x} = h(x) + wx + I(t) \tag{2.3.1}$$

where $w$ is the strength of the autapse and does not change over time. We assume a linear autapse for simplicity. The Jacobian of this system is

$$J = \frac{\partial \dot{x}}{\partial x} = \frac{1}{\tau}\Big[\frac{\partial h}{\partial x} + w\Big] \tag{2.3.2}$$

and by substituting in assumption (2.2.4), we see that the Jacobian $J$ is upper-bounded as follows:

$$J \leq \frac{1}{\tau}\Big[-\beta + w\Big] \tag{2.3.3}$$

from which we may conclude that if the following relationship holds between $w$ and $\beta$:

$$w + \beta \leq -c, \quad c > 0 \tag{2.3.4}$$

the Jacobian of this single neuron system is negative definite, and therefore the system is contracting, with rate $\frac{c}{\tau}$.

#### 2.4 Leaky Neuron with a Dynamic Autapse

What happens if $w$ is not static? In particular, if $w$ is allowed to change itself as a function of the current $w$ and $x$? Then our system becomes two-dimensional instead of one-dimensional, and finding sufficient conditions for contraction in this system becomes nontrivial. As a particular example (and one that will become useful later on), let's assume that $w$ follows the anti-hebbian dynamics

$$\dot{w} = -\gamma w - kx^2 \tag{2.4.1}$$

where $\gamma > 0$ controls the decay of forgetting and $k > 0$ controls the rate of learning in the autapse. Assume that the neuron follows the same dynamics as above, defined by (2.3.1). We now define the state vector of the whole system, $\mathbf{z}$, which is two dimensional. One dimension is the neural activation and the other dimension is the synaptic activation:

$$\mathbf{z} = \begin{bmatrix} x \\ w \end{bmatrix}$$

The state of the system at any time $t$ is a point in this $(x, w)$ space. As the system evolves, this point moves around and traces out a trajectory in this space. Of course, contraction of this state vector implies contraction of its individual elements. The dynamics of $\mathbf{z}$ are:

$$\dot{\mathbf{z}} = \begin{bmatrix} \dot{x} \\ \dot{w} \end{bmatrix} = \begin{bmatrix} h(x) + wx \\ -\gamma w - kx^2 \end{bmatrix} + \begin{bmatrix} I(t) \\ 0 \end{bmatrix}$$

Calculated the Jacobian of this system, we see that

$$\mathbf{J} = \begin{bmatrix} \frac{\partial \dot{x}}{\partial x} & \frac{\partial \dot{x}}{\partial w} \\ \frac{\partial \dot{w}}{\partial x} & \frac{\partial \dot{w}}{\partial w} \end{bmatrix} = \begin{bmatrix} \frac{\partial h}{\partial x} + w(t) & x \\ -2kx & -\gamma \end{bmatrix}$$

To conclude contraction of $\mathbf{z}$, and therefore contraction of the neural and synaptic activations, we need to show that the generalized Jacobian–see (1.2.3)–is negative definite. We begin by turning our attention to the off-diagonal elements of $\mathbf{J}$. Consider the metric defined by

$$\mathbf{\Theta} = \begin{bmatrix} 1 & 0 \\ 0 & \frac{1}{\sqrt{2k}} \end{bmatrix} \tag{2.4.2}$$

Which yields a generalized Jacobian

$$\mathbf{F} = \mathbf{\Theta J \Theta}^{-1} = \begin{bmatrix} \frac{\partial h}{\partial x} + w(t) & \sqrt{2k}x \\ -\sqrt{2k}x & -\gamma \end{bmatrix} \tag{2.4.3}$$

To show that this system is contracting, we use $\mu_2$ (the maximum eigenvalue of $\mathbf{F}_s$). We first compute $\mathbf{F}_s$ and find that

$$\mathbf{F}_s = \frac{1}{2}\Big(\mathbf{F} + \mathbf{F}^T\Big) = \begin{bmatrix} \frac{\partial h}{\partial x} + w(t) & 0 \\ 0 & -\gamma \end{bmatrix} \tag{2.4.4}$$

Since the eigenvalues of $\mathbf{F}_s$ lie along its diagonal , we need to first show that (2.3.4) is true after some finite time. To show that (2.3.4) is true, first note that the dynamics of $w$ obey the inequality

$$\dot{w} = -\gamma w - kx^2 \leq -\gamma w$$

where we have used the fact that $kx^2 \geq 0$. This implies that any *solution* of $w(t)$ obeys the inequality

$$w(t) \leq w(0)e^{-\gamma t}$$

So regardless of its initial value, $w$ will satisfy (2.3.4) exponentially. In particular, assuming that $w(0) > \beta$, the time it takes to satisfy (2.3.4), which we call $t_\beta$, is upper-bounded as follows:

$$t_\beta \leq \frac{1}{\gamma} \ln\Big(\frac{w(0)}{\beta}\Big)$$

Moreover, it will imply that the autapse in this single neuron system will become inhibitory. This makes intuitive sense–an excitatory autapse will quickly lead to runaway excitation (and thus not contraction) in the neuron. Returning to (2.4.4) and recalling that $-\gamma < 0$ by assumption, we find that

$$\mu_2(\mathbf{F}) = -\lambda = \min\Big\{|\frac{\partial h}{\partial x} + w(t)|, \gamma\Big\} < 0$$

and thus this single neuron, single synapse system is globally contracting (after exponential transients) with rate $\lambda$. Later on we will perform a similar analysis for an RNN of arbitrary size and anti-hebbian synaptic dynamics.

### 3 Synaptic Dynamics that Favor or Preserve Contraction

The assumption that synaptic dynamics are local introduces strong structure into the Jacobian of the overall network. In particular, it implies that the 'synaptic' diagonal block of the Jacobian will be a diagonal matrix. We'll now establish some simple facts about matrices with this structure that will be useful later on.

**Theorem 1.** *Consider the following matrix*

$$\mathbf{F} = \begin{bmatrix} \mathbf{X} & \mathbf{GA} \\ -\mathbf{BG}^T & \mathbf{D} \end{bmatrix}$$

*where* $\mathbf{X}$ *is an arbitrary negative definite matrix,* $\mathbf{D}$ *is diagonal and negative definite,* $\mathbf{A}$ *and* $\mathbf{B}$ *are diagonal and positive definite and* $\mathbf{G}$ *is arbitrary. Then there exists a diagonal metric* $\mathbf{\Theta}$ *such that*

$$\mathbf{\Theta F \Theta}^{-1} \prec 0 \tag{3.0.1}$$

*Proof.* Consider the candidate metric

$$\mathbf{\Theta} = \begin{bmatrix} \mathbf{I} & \mathbf{0} \\ \mathbf{0} & \mathbf{L} \end{bmatrix}$$

where $\mathbf{L}$ is some nonsingular matrix, to be determined. Applying this to the matrix $\mathbf{F}$, we find that

$$\mathbf{\Theta F \Theta}^{-1} = \begin{bmatrix} \mathbf{X} & \mathbf{GAL}^{-1} \\ -\mathbf{LBG}^T & \mathbf{D} \end{bmatrix}$$

We would like the off-diagonal terms of this matrix to cancel out when taking the symmetric part. To make this happen, we need

$$\mathbf{BL} = \mathbf{AL}^{-1}$$

Which can be easily manipulated into

$$\mathbf{L}^2 = \mathbf{AB}^{-1}$$

and thus

$$\mathbf{L} = \sqrt{\mathbf{AB}^{-1}}$$

The square root is guaranteed to exist because **A** and **B** are both positive definite and diagonal by assumption. Taking the symmetric part of (3.0.1), we find that:

$$\mathbf{\Theta F \Theta}^{-1} + (\mathbf{\Theta F \Theta}^{-1})^T = \begin{bmatrix} \mathbf{X} + \mathbf{X}^T & 0 \\ 0 & 2\mathbf{D} \end{bmatrix}$$

which is negative definite since the diagonal blocks are negative definite by assumption. This proves the result.

□

We will use this result in the next section to show that anti-hebbian plasticity gives rise to contracting dynamics.

#### 3.1 Anti-Hebbian Dynamics Produce Contraction

Consider RNNs of the following form:

$$\dot{\mathbf{x}} = \mathbf{f}(\mathbf{x}, \mathbf{W}, t) = \mathbf{h}(\mathbf{x}) + \mathbf{Wx} + \mathbf{u}(t)$$

Where $h_i(x_i)$ is a leak-term (see (2.2.4) for definition), **W** is the weight matrix of the network, and $\mathbf{u}(t)$ is a time-varying input. And now consider the anti-Hebbian dynamics:

$$\forall i \neq j \;\; \dot{w}_{ij} = -x_i x_j - \gamma(t) w_{ij} \;\text{ and }\; \dot{w}_{ii} = -\frac{1}{2}(x_i)^2 - \gamma(t) w_{ii} \tag{3.1.1}$$

with $\gamma(t) > 0$. The reason for the $\frac{1}{2}$ will be clear shortly. We can write the synaptic dynamics in an $N^2 \times 1$ vector generically as:

$$\dot{\mathbf{v}}_W = \mathbf{g}(\mathbf{v}_W, \mathbf{x})$$

where $\mathbf{v}_W$ denotes the vectorization of $\mathbf{W}$. The Jacobian of the overall neural-synaptic system is then:

$$\mathbf{J} = \begin{bmatrix} \frac{\partial \mathbf{f}}{\partial \mathbf{x}} & \frac{\partial \mathbf{f}}{\partial \mathbf{v_W}} \\ \frac{\partial \mathbf{g}}{\partial \mathbf{x}} & \frac{\partial \mathbf{g}}{\partial \mathbf{v_W}} \end{bmatrix} \tag{3.1.2}$$

##### 3.1.1 Off-Diagonal Blocks of the Jacobian

First we will show that $w_{ij}$ converges to $w_{ji}$ after exponential transients. Consider a simple distance function between $w_{ij}$ and $w_{ji}$:

$$V = (w_{ij} - w_{ji})^2$$

evaluating the time-derivative of $V$ and substituting in (3.1.1), we see that

$$\dot{V} = 2(w_{ij} - w_{ji})(-x_i x_j - \gamma(t) w_{ij} + x_i x_j + \gamma(t) w_{ji}) = -2\gamma(t) V$$

which implies that the distance between $w_{ij}$ and $w_{ji}$ shrinks exponentially with at least rate $\min_t(\gamma(t))$, independently of initial condition or the particular trajectory of $\mathbf{x}$. For the rest of the analysis we will therefore assume that $w_{ij} \approx w_{ji}$. Using the symmetry of $\mathbf{W}$, the following four relationship can be shown:

$$\forall i \neq j \;\; \frac{\partial \dot{w}_{ij}}{\partial x_i} = -x_j \;\text{ and that }\; \frac{\partial \dot{x}_i}{\partial w_{ij}} = x_j$$

as well as

$$\forall i \neq j \ \ \frac{\partial \dot{w}_{ji}}{\partial x_i} = -x_j \ \text{ and that } \ \frac{\partial \dot{x}_i}{\partial w_{ji}} = x_j$$

The first three of these relationships can be verified by direct differentiation of (3.1.1). The last relationship can be seen by using the symmetry of $\mathbf{W}$. In particular, that:

$$\frac{\partial \dot{x}_i}{\partial w_{ji}} \approx \frac{\partial \dot{x}_i}{\partial w_{ij}} = x_j$$

What we have shown now is that under the synaptic dynamics defined above, we have the following relationship:

$$\frac{\partial \mathbf{g}}{\partial \mathbf{x}} = -\frac{\partial \mathbf{f}}{\partial \mathbf{v}_W}^T \tag{3.1.3}$$

Therefore, the off-diagonal blocks in the overall system Jacobian (3.1.2) will cancel out when computing its symmetric part.

##### 3.1.2 Diagonal Blocks of the Jacobian

The upper left block of the Jacobian is

$$\frac{\partial \mathbf{f}}{\partial \mathbf{x}} = \frac{\partial \mathbf{h}}{\partial \mathbf{x}} + \mathbf{W}(t)$$

thus, for this matrix to be negative definite we need

$$\mu_2(\mathbf{W}(t)) < \beta \tag{3.1.4}$$

To see that this will be the case, consider that the synaptic dynamics defined by (3.1.1) can be written in matrix notation as

$$\dot{\mathbf{W}} = -\mathbf{P} \odot \mathbf{H} - \gamma(t)\mathbf{W}$$

where

$$P_{ij} = \begin{cases} 1 & \text{if } i \neq j \\ \frac{1}{2} & \text{if } i = j \end{cases}$$

Now discretize the differential equations for $\mathbf{W}$:

$$\dot{\mathbf{W}} \approx \frac{\mathbf{W}_{t+1} - \mathbf{W}_t}{\Delta t} = -\mathbf{P} \odot \mathbf{x}_t \mathbf{x}_t^T - \mathbf{W}_t$$

rearranging this to collect all the $t$ terms on the right hand side, we have

$$\mathbf{W}_{t+1} = (1 - \Delta t)\mathbf{W}_t - \Delta t \mathbf{P} \odot \mathbf{x}_t \mathbf{x}_t^T$$

To make derivations less tedious, let's define $\beta \equiv \Delta t > 0$ and $\alpha \equiv (1 - \beta)$. Note that since that, ideally, $\Delta t$ is small, we can say that $0 < \alpha < 1$. Now we have the discrete dynamical system for $\mathbf{W}$:

$$\mathbf{W}_{t+1} = \alpha \mathbf{W}_t - \beta \mathbf{P} \odot \mathbf{x}_t \mathbf{x}_t^T$$

We know need to notice that $-\mathbf{P} \odot \mathbf{x}_t \mathbf{x}_t^T$ is symmetric (since $\mathbf{P}$ and $\mathbf{x}_t \mathbf{x}_t^T$ are symmetric). It is also negative semi-definite, by the Schur Product Theorem. Let's denote this expression by $\mathbf{N}_t = \mathbf{N}_t^T \prec 0$ so that

$$\mathbf{W}_{t+1} = \alpha \mathbf{W}_t + \beta \mathbf{N}_t$$

Taking the symmetric part of both sides we find that:

$$(\mathbf{W}_{t+1})_s = \alpha (\mathbf{W}_t)_s + \beta \mathbf{N}_t$$

We can now use Weyl's inequality for the sum of Hermitian matrices (see premilinaries) to conclude that

$$\mu_2(\mathbf{W}_{t+1}) \leq \alpha \mu_2(\mathbf{W}_t)$$

and therefore that

$$\mu_2(\mathbf{W}_t)) \leq \alpha^t \mu_2(\mathbf{W}_0)$$

which, since $0 < \alpha < 1$, means that the symmetric part of $\mathbf{W}$ decays exponentially to zero. Recall that for contraction we needed $\mu_2(\mathbf{W}) < \beta$. Assuming that $\mu_2(\mathbf{W}_0) > \beta$, the upper-bound for the number of steps this will take is $O(\ln(\frac{\mu_2(\mathbf{W}_0)}{\beta}))$. See the figure below for a simulation confirming this result.

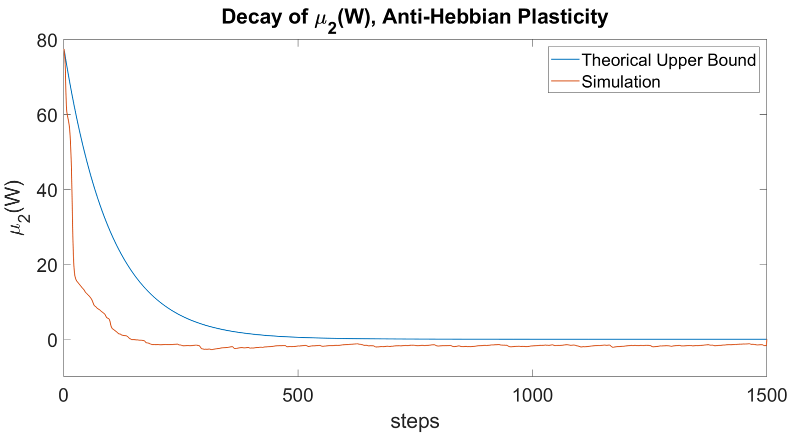

#### 3.2 Generalized to Different Anti-Hebbian Gains

We can also use (3.1.3) to generalize the above result to different $\mathbf{P}$ matrices. That is, to different anti-Hebbian gains. To do so let's first define the 'Hebbian' term as $\mathbf{H} \equiv \mathbf{x}\mathbf{x}^T$ Now the synaptic dynamics defined by (3.1.1) can in matrix notation can be written as

$$\dot{\mathbf{W}} = -\mathbf{P} \odot \mathbf{H} - \gamma(t)\mathbf{W} \tag{3.2.1}$$

where

$$P_{ij} = \begin{cases} 1 & \text{if } i \neq j \\ \frac{1}{2} & \text{if } i = j \end{cases}$$

vectorizing (3.2.1) and using the relation (1.1.11), we can see that

$$\dot{\mathbf{v}}_W = -\mathbf{D}_P \mathbf{v}_H - \gamma(t)\mathbf{v}_W$$

and differentiating this expression with respect to $\mathbf{x}$, we have:

$$\frac{\partial \dot{\mathbf{v}}_W}{\partial \mathbf{x}} = -\mathbf{D}_P \frac{\partial \mathbf{v}_H}{\partial \mathbf{x}} = -\frac{\partial \mathbf{f}}{\partial \mathbf{v}_W}$$

and finally, left multiplying this expression by the inverse of $\mathbf{D}_P$ we have

$$\frac{\partial \mathbf{v}_H}{\partial \mathbf{x}} = \mathbf{D}_P^{-1} \frac{\partial \mathbf{f}}{\partial \mathbf{w}}^T \tag{3.2.2}$$

Note that this expression is only true when $\mathbf{W}$ is symmetric. Now imagine we have synaptic dynamics of the more general form:

$$\dot{\mathbf{W}} = -\mathbf{K} \odot \mathbf{H} - \gamma(t)\mathbf{W}$$

where $\mathbf{K}$ is symmetric (to preserve to convergence of $\mathbf{W}$ to a symmetric matrix), positive semi-definite (to preserve negative-definiteness of the upper right Jacobian block) and has positive entries (for reasons explained right now). Vectorizing in the same manner as above and substituting in (3.2.2), we find that

$$\frac{\partial \mathbf{v}_{\dot{W}}}{\partial \mathbf{x}} = -\mathbf{D}_K \frac{\partial \mathbf{v}_H}{\partial \mathbf{x}} = -\mathbf{D}_K \mathbf{D}_P^{-1} \frac{\partial \mathbf{f}}{\partial \mathbf{v}_W}$$

If $\mathbf{D}_K$ is positive (which will happen when $\mathbf{K}$ has all positive entries), then $\mathbf{D}_K \mathbf{D}_P^{-1}$ is diagonal and positive definite, and thus Theorem 1 applies–this is a feedback combination that automatically preserves contraction.

To summarize, we have shown that the synaptic dynamics described by

$$\dot{\mathbf{W}} = -\mathbf{K} \odot \mathbf{x}\mathbf{x}^T - \gamma(t)\mathbf{W} \tag{3.2.3}$$

where $\mathbf{K}$ is a symmetric, positive semi-definite matrix with positive entries, lead to contracting neural and synaptic dynamics in a wide class of RNNs. We did this by showing that the overall Jacobian (3.1.2) has diagonal blocks that 'cancel' out and diagonal blocks that are negative definite–implying negativedefinitness of the whole Jacobian and thus contraction of the combined neural and synaptic system.

### 4 Sparsity Ensures Contraction

In this section we show that it is possible to ensure contraction–without any knowledge of the specific synaptic dynamics–by keeping the network sufficiently sparse. This result is a straightfoward application of the $\mu_\infty$ matrix measure to the Jacobian of the overall neural and synaptic system. Consider neural networks of the form

$$\dot{x}_i = h(x_i) + \sum_{j \neq i}^{N} W_{ij} r_j + u_i(t) \tag{4.0.1}$$

where $h_i$ is a leak-term and $r_i$ is bounded activation function that satisfies the following two properties:

$$|r_i| \leq r_{max} \text{ and } 0 < g_{min} \leq \frac{\partial r_i}{\partial x_i} \leq g_{max} \tag{4.0.2}$$

We consider synaptic dynamics of the form

$$\dot{W}_{ij} = g(W_{ij}, x_i, x_j) \tag{4.0.3}$$

That is to say, we only consider synaptic dynamics that are *local*, in the sense that the change in weight is only a function of the weight itself, and the pre and post synaptic neurons.

It will be useful to define, for neuron $i$, the total number of afferent synapses and the total number of afferent *dynamic* synapses which we denote $p_i$ and $d_i$, respectively. Note that $d_i$ is a fraction of $p_i$ and can therefore be written as $d_i = \alpha_i p_i$ for some $0 \leq \alpha_i \leq 1$. It will also be useful to give names to one more global system parameter: the maximum amplitude (absolute value) of all the weights in the network: $w_* \equiv \max_{ij}(|w_{ij}|)$. For the neural rows of the Jacobian, we see that the $\mu_\infty$ measure can be upper bounded as

$$sup_i(\frac{\partial h_i}{\partial x_i} + \sum_{j \neq i}^{N} |\frac{\partial \dot{x}_i}{\partial x_i}| + \sum_{q=1}^{d_i} |\frac{\partial \dot{x}_i}{\partial W_{iq}}|) < 0 \tag{4.0.4}$$

By using the following two relations:

$$\frac{\partial \dot{x}_i}{\partial x_j} = W_{ij} \frac{\partial r_j}{\partial x_j} \text{ and } \frac{\partial \dot{x}_i}{\partial W_{ij}} = r_j$$

we can see that (4.0.4) can be assured if

$$\forall i, t \;\; p_i g_{max} w_* + d_i r_{max} < \beta$$

Substituting in the relationship $d_i = \alpha_i p_i$ into the above, we get:

$$\forall i, t \;\; p_i[g_{max} w_* + \alpha_i r_{max}] < \beta \tag{4.0.5}$$

We still have the 'synaptic' rows of the Jacobian to worry about, but these are much simpler. The locality assumption implies that there are *at most* three elements per row. These elements are the derivatives of $g_{ij}$ with respect to $x_i$ and $x_j$ and the derivative of $g_{ij}$ with respect to $w_{ij}$. Thus, if

$$\forall i, j, t \;\; |\frac{\partial g_{ij}}{\partial x_i}| + |\frac{\partial g_{ij}}{\partial x_j}| + \frac{\partial g_{ij}}{\partial w_{ij}} < 0 \tag{4.0.6}$$

the bottom two Jacobian blocks satisfy the matrix measure condition. Intuitively, this inequality tells us that the 'self-stabilization' of the synapses must be stronger than the change in plasticity due to neural activity. For example, if one imagines that $g_{ij}$ contains a constant leak term of the form $-\gamma w_{ij}$ where $\gamma > 0$, then the inequality above is a sufficient condition on how large $\gamma$ should be.

In summary, we have shown that if the synapses are contracting (i.e (4.0.6) is satisfied) then (4.0.5) tells us that it is possible to assure contraction of the overall neural and synaptic network by making the network sufficiently sparse.

### 5 Contraction in RNNs with Static Weights

In this section we'll analyze RNNs of the general form

$$\dot{x}_i = h_i(x_i) + \sum_{i=1}^{n} W_{ij}\phi(g_i x_i) + u_i(t) \tag{5.0.1}$$

Where $h_i$ is a leak-term (see (2.2.4) for definition), $\phi$ is a nonlinear function with a linear 'regime' (such as tanh(x)), $g_i$ is a neuron-specific gain-term that satisfies:

$$\forall i, \;\; 0 < g_{min} \leq g_i \leq g_{max}$$

and $u_i(t)$ is a time-varying input into neuron $i$. For simplicity, we focus here on the regime where

$$\phi(g_i x_i) \approx g_i x_i$$

In this case we may rewrite (5.0.1) in matrix form as:

$$\dot{\mathbf{x}} = \mathbf{h}(\mathbf{x}) + \mathbf{W}\mathbf{G}\mathbf{x} + \mathbf{u}(t) \tag{5.0.2}$$

where $\mathbf{G}$ is a constant diagonal, positive definite matrix such that $G_{ii} = g_i$. The Jacobian of (5.0.1) is a state-dependent matrix:

$$\mathbf{J} = \mathbf{L} + \mathbf{W}\mathbf{G} \tag{5.0.3}$$

where $\mathbf{L}$ is a diagonal matrix such that $L_{ii} = \frac{\partial h_i}{\partial x_i} \leq -\beta$. Consider the following metric transformation to yield a generalized Jacobian, $\mathbf{F}$:

$$\mathbf{\Theta} = \mathbf{G}^{1/2}\mathbf{D} \tag{5.0.4}$$

Where $\mathbf{D}$ is an arbitrary, non-singular diagonal matrix. Applying this metric to (5.0.3) we find that

$$\begin{aligned} \mathbf{F} &= \mathbf{\Theta}\mathbf{J}\mathbf{\Theta}^{-1} \\ &= \mathbf{\Theta}\mathbf{L}\mathbf{\Theta}^{-1} + \mathbf{\Theta}\mathbf{W}\mathbf{G}\mathbf{\Theta}^{-1} \\ &= \mathbf{L} + \mathbf{G}^{1/2}[\mathbf{D}\mathbf{W}\mathbf{D}^{-1}]\mathbf{G}^{1/2} \\ &= \mathbf{G}^{1/2}[\mathbf{G}^{-1}\mathbf{L} + \mathbf{D}\mathbf{W}\mathbf{D}^{-1}]\mathbf{G}^{1/2} \end{aligned} \tag{5.0.5}$$

Which implies that a sufficient condition for contraction in the RNN described by (5.0.2) is that

$$\mu_{D,2}(\mathbf{W}) + \frac{\beta}{g_{max}} \leq -c, \quad c > 0 \tag{5.0.6}$$

If (5.0.6) is true, then the network described by (5.0.2) is contracting with rate of $\lambda_F = c\, g_{min}$.

##### 5.0.1 Contraction in Circuits with Perfect E-I Balance

One practical implication of (5.0.6) is that we can now exploit feedback structure present in $\mathbf{W}$ in a way that would have been missed by using the spectral norm. As an example, consider a network with weight matrix

$$\mathbf{W} = \begin{bmatrix} \mathbf{W}_{EE} & \mathbf{W}_{EI} \\ \mathbf{W}_{IE} & \mathbf{W}_{II} \end{bmatrix} \tag{5.0.7}$$

Assume that $\mathbf{W}_{IE}$ is a matrix containing all positive entries and $\mathbf{W}_{EI}$ is a matrix containing all negative entries. Further assume that:

$$-k\mathbf{W}_{IE}^T = \mathbf{W}_{EI} \tag{5.0.8}$$

where $k > 0$. In other words, (5.0.8) asserts that this circuit has perfect E-I balance, up to a constant scalar factor. In this case, the weight matrix becomes:

$$\mathbf{W} = \begin{bmatrix} \mathbf{W}_{EE} & -k\mathbf{W}_{IE}^T \\ \mathbf{W}_{IE} & \mathbf{W}_{II} \end{bmatrix}$$

This corresponds to two interacting neural populations, one inhibitory and one excitatory. The excitatory population excites the inhibitory one through the $\mathbf{W}_{IE}$ submatrix and the inhibitory population inhibits the excitatory population through the $-k\mathbf{W}_{IE}^T$ submatrix. Denote the (unweighted) matrix measures of the EE and II subnetworks $\mu_{EE}$ and $\mu_{II}$, respectively:

$$\mu_2(\mathbf{W}_{EE}) = \mu_{EE} \ \text{ and } \ \mu_2(\mathbf{W}_{II}) = \mu_{II} \tag{5.0.9}$$

It's straightforward to show that by picking $\mathbf{D}$ as:

$$\mathbf{D} = \begin{bmatrix} \mathbf{I} & 0 \\ 0 & \sqrt{k}\mathbf{I} \end{bmatrix}$$

we get:

$$\mu_{D,2}(\mathbf{W}) = \frac{1}{2}\lambda_{\max}([\mathbf{DWD}^{-1} + \mathbf{D}^{-1}\mathbf{W}^T\mathbf{D}])$$
$$= \frac{1}{2}\lambda_{\max}(\begin{bmatrix} \mathbf{W}_{EE} + \mathbf{W}_{EE}^T & 0 \\ 0 & \mathbf{W}_{II} + \mathbf{W}_{II}^T \end{bmatrix}) \tag{5.0.10}$$
$$= \max[\mu_2(\mathbf{W}_{EE}), \mu_2(\mathbf{W}_{II})]$$

which implies that

$$\mu_{D,2}(\mathbf{W}) = \max[\mu_{EE}, \mu_{II}]$$

This shows that by appropriately scaling $\mu_{EE}$ and $\mu_{II}$ to satisfy (5.0.6), the RNN can be made contracting *independent* of the strength of individual elements in the diagonal blocks.

##### 5.0.2 Contraction in Circuits With Statistical E-I Balance

In a biological system, perfect E-I balance (i.e (5.0.8)) cannot reasonably be expected. However, some basic results from matrix algebra and random matrix theory can tell us that contraction can be assured even in the case where E-I balance occurs in an average sense.

In particular, we'll need the following result from matrix algebra. Consider a symmetric block matrix of the form:

$$\begin{bmatrix} \mathbf{A} & \mathbf{B} \\ \mathbf{B}^T & \mathbf{C} \end{bmatrix} \tag{5.0.11}$$

where $\mathbf{A}$ and $\mathbf{C}$ are negative definite and symmetric, with $\lambda_{\max}(\mathbf{A}) = -\lambda_A < 0$ and $\lambda_{\max}(\mathbf{C}) = -\lambda_C < 0$. Then a standard result from matrix algebra [3] says that a sufficient condition for negative-definiteness of (5.0.11) is that:

$$||\mathbf{B}||_2^2 < \lambda_A \lambda_C \tag{5.0.12}$$

We'll also need the following result from random matrix theory. Consider an $N \times N$ symmetric matrix, $\mathbf{R}$, whose elements above and including the diagonal are distributed according to a Gaussian distribution:

$$R_{ij} = \mathcal{N}(0, \sigma^2) \text{ for } j \geq i$$

Then for large $N$, we have from [2] that, in expectation:

$$||\mathbf{R}||_2 \approx 2\sigma\sqrt{N} \tag{5.0.13}$$

We can use (5.0.12) together with (5.0.13) for how much deviation from perfect E-I balance a circuit of size $N$ can tolerate while preserving contraction. In particular, consider again the matrix defined in (5.0.7) and assume that (5.0.6) is satisfied for each diagonal block. That is, assume:

$$\mu_{EE} + \beta g_{max}^{-1} \leq -c \text{ and } \mu_{EE} + \beta g_{max}^{-1} \leq -c \text{ with } c > 0$$

and assume $N_E = N_I = \frac{N}{2}$ for simplicity. We define $\mathbf{R}$ as follows:

$$\mathbf{R} \equiv \frac{\mathbf{W}_{EI} + \mathbf{W}_{IE}^T}{2}$$

from (5.0.12), we can assure contraction in regions where:

$$||\mathbf{R}||_2 < c$$

To find this region, we will upper bound the spectral norm of $\mathbf{R}$ as follows: instead of the case where E-I and I-E connections cancel out exactly, consider the case when instead they are each drawn from a distribution whose *means* cancel out exactly. Namely:

$$W_{IE}^{ij} \sim \mathcal{N}(m, \sigma_{IE}^2) \text{ and } W_{EI}^{ij} \sim \mathcal{N}(-m, \sigma_{EI}^2)$$

This implies that:

$$R_{ij} = \frac{W_{IE}^{ij} + W_{EI}^{ji}}{2} \sim \mathcal{N}(0, \frac{\sigma_{IE}^2 + \sigma_{EI}^2}{4})$$

Thus, the matrix $\mathbf{R}$ is a symmetric, Gaussian matrix with zero mean and variance $\frac{\sigma_{IE}^2 + \sigma_{EI}^2}{4}$. By plugging this expression into (5.0.13) we can conclude contraction when the following relationship holds between $\mathbf{R}$ and the eigenvalues of the E-E and I-I connection matrices:

$$\sigma_{IE}^2 + \sigma_{EI}^2 < \frac{c^2}{N}$$

Of course, in the limit as the spreads go to zero, we arrive back at the case of perfect E-I balance with $k = 1$, which always yields contraction.

Another way to get a sense of the tolerance' for imperfect E-I balance is to use the Frobenius norm of $\mathbf{R}$, which upper-bounds the spectral norm. Assume that for each connection between excitatory neuron $i$ and inhibitory neuron $j$, we have that

$$|W_{EI}^{ij} + W_{IE}^{ji}|^2 \leq \kappa^2 \text{ with } \kappa > 0$$

This straightforwardly implies that the Frobenius norm of $\mathbf{R}$ is upper bounded by:

$$||\mathbf{R}||_F \leq \kappa N$$

Which means that it is possible to ensure contraction by scaling $\kappa$ with $N$, since:

$$||\mathbf{R}||_2 \leq ||\mathbf{R}||_F \leq \kappa N < c$$

So by scaling $\kappa$, which measures how imperfectly each E-I synapse is balanced by the corresponding I-E synapse, we can guarantee contraction.

#### 5.1 Relationship to Echo State Networks

Echo state networks (ESNs) are contracting, nonlinear, discrete-time RNNs with static synaptic weights. While attempts have been made to apply the Echo-State framework to continuous time systems, these attempts have relied on discretizations of the underlying continuous dynamics. What determines if a network is an ESN is the 'Echo-State Property' (ESP), which is typically in the form of a bound on the spectral norm of the weight matrix. Unlike our result above (see 5.0.6) this does not allow one to factor in stabilizing influences within the weight matrix, such as E-I balance. Nevertheless, contraction analysis can be applied to discrete time systems. For completeness, we will now do this to and derive a slight generalization of a known sufficient condition for the ESP in networks where the activation function is more flexible than $\tanh(x)$.

We will need the following result. Consider a discrete-time dynamical system:

$$\mathbf{x}_{n+1} = \mathbf{T}(\mathbf{x}_n, \mathbf{u}_n)$$

It was shown in [4] that this system will be contracting if there exists some non-singular matrix $\mathbf{\Theta}_n$ such that

$$\sigma_{\max}(\mathbf{\Theta}_{n+1}\frac{\partial \mathbf{T}}{\partial \mathbf{x}}\mathbf{\Theta}_n^{-1}) < 1$$

**Theorem 2.** *Consider the discrete time RNN*

$$\mathbf{x}_{n+1} = \mathbf{W}\mathbf{r}(\mathbf{x}_n + \mathbf{u}_n) \tag{5.1.1}$$

*where $r_i = r_j = r$ is a nonlinearity (e.g. tanh(x)) that satisfies*

$$0 \leq \frac{\partial r}{\partial x}(x) \leq g \tag{5.1.2}$$

*If the matrix $\tilde{\mathbf{W}} \equiv \sqrt{g}\mathbf{W}$ is Schur diagonally stable, then the RNN is contracting, and therefore has the ESP. That is, if there exists a postive definite, diagonal matrix $\mathbf{P}$ such that:*

$$\tilde{\mathbf{W}}^T\mathbf{P}\tilde{\mathbf{W}} - \mathbf{P} < 0 \;\; \textit{with} \;\; \tilde{\mathbf{W}} \equiv \sqrt{g}\mathbf{W}$$

*the RNN is contracting.*

*Proof.* For the RNN defined in the theorem, we have

$$\sigma_{\max}(\mathbf{\Theta}_{n+1}\mathbf{W}\mathbf{\Phi}\mathbf{\Theta}_n^{-1}) < 1$$

Let $\Theta_n = \mathbf{P}^{1/2}$, where $\mathbf{P}$ is diagonal and positive definite. Using the submultiplicitivity of the spectral norm, we find that a sufficient condition for contraction is:

$$\sigma_{\max}(\mathbf{P}^{1/2}\mathbf{W}\mathbf{P}^{-1/2}) < g^{-1}$$

we can easily rewrite this condition as

$$\mathbf{W}^T\mathbf{P}\mathbf{W} - \mathbf{P}g^{-1} < 0$$

Multiplying both sides by $g$, we find that the RNN is contracting if

$$g\mathbf{W}^T\mathbf{P}\mathbf{W} - \mathbf{P} < 0 \iff \tilde{\mathbf{W}}^T\mathbf{P}\tilde{\mathbf{W}} - \mathbf{P} < 0$$

Which can be readily recognized as the statement that the matrix $\tilde{\mathbf{W}} = \sqrt{g}\mathbf{W}$ is Schur diagonally stable. □

**Remark 1.** *The above stability criterion immediatly applies to RNNs of the form:*

$$\mathbf{x}_{n+1} = \mathbf{r}(\mathbf{W}\mathbf{x}_n + \mathbf{u}_n)$$

*for nonsingular* $\mathbf{W}$*. This can be seen using the following coordinate transform and applying it to* (5.1.1)*:*

$$\mathbf{y} = \mathbf{W}^{-1}\mathbf{x}$$

$$\mathbf{y}_{n+1} = \mathbf{W}^{-1}\mathbf{x}_{n+1} = \mathbf{W}^{-1}[\mathbf{W}\mathbf{r}(\mathbf{x}_n + \mathbf{u}_n)] = \mathbf{r}(\mathbf{W}\mathbf{y}_n + \mathbf{u}_n)$$

*Since* $\delta\mathbf{y} = \mathbf{W}^{-1}\delta\mathbf{x}$*, this means that* $\delta\mathbf{x} \to 0 \iff \delta\mathbf{y} \to 0$*,*

**Remark 2.** *For the specific case of* $r(x) = \tanh(x)$*,and therefore* $g = 1$*, this result already appears in [8].*

**Remark 3.** *A common RNN uses the discontinuous activation function:*

$$\mathbf{r}(x_i) = \max(0, x_i)$$

*This does not pose a problem for the above stability condition, for the following reason. Define a diagonal, 'switching' matrix* $\mathbf{S}$ *in the following way:*

$$S_{ii} = \begin{cases} 1 & \text{if } x_i > 0 \\ 0 & \text{otherwise} \end{cases}$$

*And note that the dynamics of the ReLu network can be written:*

$$\mathbf{x}_{n+1} = \mathbf{W}\mathbf{S}\mathbf{x}_n + \mathbf{u}_n$$

*The Jacobian of this system is:*

$$\mathbf{J} = \mathbf{W}\mathbf{S}$$

*Since* $\mathbf{S}$ *is diagonal and* $||\mathbf{S}|| \leq 1$*, we can use the same diagonal metric transformation as above to find that a sufficient condition for a ReLu RNN to be contracting is that* $\mathbf{W}$ *is Schur diagonally stable. In other words, if there exists some positive and diagonal* $\mathbf{P}$ *such that*

$$\mathbf{W}^T\mathbf{P}\mathbf{W} - \mathbf{P} < 0$$

*the ReLu network is contracting.*

### 6 Setting Neural Operating Points

Consider the following autonomous system, contracting with rate $\lambda$ in metric $\mathbf{\Theta}$:

$$\dot{\mathbf{x}} = \mathbf{f}(\mathbf{x}) + \mathbf{c}$$

Because this system is autonomous and contracting, it will converge exponentially to a fixed point $\mathbf{x}^*$ which is the unique solution to:

$$\mathbf{f}(\mathbf{x}^*) = -\mathbf{c}$$

Now add an arbitrary time-varying input $\mathbf{u}(t)$ to the system, so that the equation reads:

$$\dot{\mathbf{y}} = \mathbf{f}(\mathbf{y}) + \mathbf{c} + \mathbf{u}(t)$$

This system will not, generally speaking, converge to a fixed point (since $\mathbf{u}(t)$ varies with time). However using the robustness results stated earlier we can bound any trajectory of $\mathbf{x}$ to $\mathbf{x}^*$. In particular, after exponential transients of rate $\lambda$ we have:

$$|\mathbf{y}(t) - \mathbf{x}^*| \leq \frac{|\mathbf{u}(t)|\chi}{\lambda}$$

where $\chi$ is an upper-bound on the condition number of $\mathbf{\Theta}$. Geometrically, this says that $\mathbf{y}(t)$ will always stay within a ball of radius $\frac{|\mathbf{u}(t)|\chi}{\lambda}$ around $\mathbf{x}^*$ (this point is the 'neural operating point' we refer to in the paper). Since $\mathbf{x}^*$ depends implicitly on $\mathbf{c}$, this means that different $\mathbf{c}$ lead to different operating points.

### 7 Metric-Induced Overshoots, Longer Information Retention, and a Hierarchy of Time Constants

At discussed earlier, metric transformations introduce a transient overshoot in exponential convergence. In particular, let $\chi$ denote an upper bound on the condition number of $\mathbf{\Theta}$. Then the distance between any two trajectories $d(t) = |\mathbf{x}(t) - \mathbf{x}'(t)|$ obeys:

$$d(t) \leq \chi e^{-\lambda t}$$

For certain feedback combinations $\chi$ can be written down analytically, and depends on basic system parameters such as coupling strength. In particular in the main text we consider coupling two linear RNNs (a PFC RNN and a sensory RNN) of the same dimension in feedback of the form

$$\dot{\mathbf{p}} = -\mathbf{p} + \mathbf{W}_p\mathbf{p} + \mathbf{G}\mathbf{s}$$

$$\dot{\mathbf{s}} = -\mathbf{s} + \mathbf{W}_s\mathbf{s} - k\mathbf{G}^T\mathbf{p}$$

where $\mathbf{G}$ is a matrix of coupling gains between the two networks and $\mathbf{W}_p$ and $\mathbf{W}_s$ contain the recurrent connections of the PFC network and sensory network, respectively. The parameter $k$ represents the strength of feedback from the PFC RNN to the sensory RNN. The reason we chose linear RNNs is because sharper contraction conditions are known for linear systems, as opposed to a general nonlinear RNN, to be contracting. This means we could bring the RNNs closer to the edge of contraction, making the network more sensitive to inputs and encouraging longer timescales–to better illustrate our point–as we combined these systems. The Jacobian of this combined system is:

$$\mathbf{J} = \begin{bmatrix} -\mathbf{I} + \mathbf{W}_p & \mathbf{G} \\ -k\mathbf{G}^T & -\mathbf{I} + \mathbf{W}_s \end{bmatrix}$$

This is the same feedback combination as we discussed earlier, so we know this system is contracting in metric:

$$\mathbf{\Theta} = \begin{bmatrix} \mathbf{I} & \mathbf{0} \\ \mathbf{0} & \frac{1}{\sqrt{k}}\mathbf{I} \end{bmatrix}$$

if $\mu_2(\mathbf{W}_p) < 1$ and $\mu_2(\mathbf{W}_p) < 1$, which is easy to ensure by generating random matrices and then scaling by the appropriate constant. For $k < 1$, we have:

$$\chi = \frac{\sigma_{\max}(\mathbf{\Theta})}{\sigma_{\min}(\mathbf{\Theta})} = \frac{1}{\sqrt{k}}$$

for $k < 1$. Thus, by decreasing $k$ towards 0, one can increase $\chi$ and thereby increase the room for overshoot and longer information retention.

We will now move on to exploring hierarchical combinations. As discussed above, hierarchical combinations (cascading combinations) of contracting systems are contracting. One can show this inductively by starting with a combination of two contracting systems and generalizing to an arbitrary number. We are interested here in explicitly deriving a metric in which the generalized Jacobian of a hierarchy of arbitrary depth is contracting. The condition number of this metric will give us an upper bound for the overshoot as a function of position in the hierarchy. This is basically an exercise in block-matrix multiplication.

Consider $L$ contracting systems, each of arbitrary dimension. Imagine that they are initially uncoupled from one another. The Jacobian of the overall system is block diagonal, where each block is Jacobian of the $L^{th}$ system with respect to itself:

$$\mathbf{J} = \begin{bmatrix} \mathbf{J}_{11} & \mathbf{0} & \dots & \mathbf{0} \\ \mathbf{0} & \mathbf{J}_{22} & \dots & \mathbf{0} \\ \vdots & \vdots & \ddots & \vdots \\ \mathbf{0} & \mathbf{0} & \dots & \mathbf{J}_{LL} \end{bmatrix}$$

Now connect the subsystems hierarchically, meaning that if subsystem A has forward connections to subsystem B, subsystem B has no foward connections to subsystem A. One can show that this induces a triangular structure on the overall Jacobian, so that it becomes:

$$\mathbf{J} = \begin{bmatrix} \mathbf{J}_{11} & \mathbf{0} & \dots & \mathbf{0} \\ \mathbf{G}_{21} & \mathbf{J}_{22} & \dots & \mathbf{0} \\ \vdots & \vdots & \ddots & \vdots \\ \mathbf{G}_{L1} & \mathbf{G}_{L2} & \dots & \mathbf{J}_{LL} \end{bmatrix}$$

where $\mathbf{G}_{ij}$ represents the coupling Jacobian between subsystems $i$ and $j$. Note that $\mathbf{G}_{ij}$ can be time-varying and state-dependent. The structure in $\mathbf{J}$ can be summed up succinctly by noting that $(i, j)^{th}$ block of $\mathbf{J}$, $\mathbf{J}_{ij}$, satisfies:

$$\mathbf{J}_{ij} = \begin{cases} \mathbf{G}_{ij} & \text{if } j \leq i \\ \mathbf{0} & \text{otherwise} \end{cases}$$

Consider the diagonal metric defined block-wise as

$$\mathbf{\Theta}_{ii} = \epsilon^{i-1}\mathbf{I}_i$$

where $\mathbf{I}_i$ is of proper dimension. It's straightfoward to show that the $(i,j)^{th}$ block of the generalized Jacobian of this system, $\mathbf{F}_{ij}$, is

$$\mathbf{F}_{ij} = \epsilon^{i-j}\mathbf{J}_{ij}$$

For $i = j$, we have that $\mathbf{F}_{ii} = \mathbf{J}_{ii}$. For $j > i$, we have that $\mathbf{F}_{ij} = \mathbf{0}$. Thus, by sending $\epsilon \to 0$ we can bring the generalized Jacobian arbitrarily close to a block diagonal matrix, and so the overall system is contracting if the individual subsystems are contracting. Assuming $\epsilon < 1$, let's write it as $\epsilon \equiv \frac{1}{x}$ for some $x > 1$. Increasing $x$ corresponding to decreasing $\epsilon$. Intuitively, the smaller $\epsilon$ is, the larger connectivity gain between subsystems is. The metric $\mathbf{\Theta}$ is now:

$$\mathbf{\Theta}_{ii} = (\frac{1}{x})^{i-1}\mathbf{I}_i = x^{1-i}\mathbf{I}_i$$

Clearly,

$$\sigma_{\max}(\mathbf{\Theta}) = 1 \text{ and } \sigma_{\min}(\mathbf{\Theta}) = x^{1-L}$$

Thus we have

$$\chi = \frac{\sigma_{\max}(\mathbf{\Theta})}{\sigma_{\min}(\mathbf{\Theta})} = x^{L-1}$$

Thus, as $x$ increases (which corresponds to stronger connectivity between subsystems), the potential overshoot grows. This makes intuitive sense, as small overshoots early on the hierarchy get multiplied through strong connections and take longer for the system to wash out. There is also the exponential scaling with the number of subsystems, which is discussed in the main text.
